# Post-activation depression produces extensor H-reflex suppression following flexor afferent conditioning

**DOI:** 10.1101/2022.03.28.486118

**Authors:** K. Metz, I. Concha-Matos, K. Hari, O. Bseis, B. Afsharipour, S. Lin, Y. Li, R. Singla, K Fenrich, DJ. Bennett, MA. Gorassini

## Abstract

Suppression of the extensor H-reflex by flexor afferent conditioning is thought to be produced by a long-lasting inhibition of extensor Ia-afferent terminals via primary afferent depolarization (PAD) activated by GABA_A_ receptors. Considering the recent finding that PAD does not produce presynaptic inhibition of Ia-afferent terminals, we examined if H-reflex suppression is instead mediated by post-activation depression of the test extensor Ia-afferents triggered by PAD-evoked spikes and/or by a long-lasting inhibition of the extensor motoneurons. A brief conditioning vibration of the flexor tendon suppressed both the extensor soleus H-reflex and the tonic discharge of soleus motor units for 300 ms, indicating that part of the H-reflex suppression was mediated by a long-lasting inhibition of the extensor motoneurons. When activating the flexor afferents electrically to produce conditioning, the soleus H-reflex was also suppressed for 300 ms, but only when a short-latency reflex was evoked in the soleus muscle by the conditioning input itself. In mice, a similar short-latency reflex was evoked when optogenetic or afferent activation of GABAergic (GAD2^+^) neurons produced PAD large enough to evoke orthodromic spikes in the test Ia-afferents, causing post-activation depression of subsequent monosynaptic excitatory-post-synaptic potentials. The time course of this post-activation depression and related H-reflex suppression (lasting 2 s) was like rate-dependent depression that is also due to post-activation depression. We conclude that extensor H-reflex inhibition by brief flexor afferent conditioning is produced by both post-activation depression of extensor Ia-afferents and long-lasting inhibition of extensor motoneurons, rather than from PAD directly inhibiting Ia afferent terminals.

## Introduction

The suppression of the H-reflex following a conditioning stimulation to an antagonist nerve was until recently thought to be mediated by the activation of a primary afferent depolarization (PAD) in the Ia afferent terminal (Hultborn *et al*., 1987a; Stein, 1995; Misiaszek, 2003; Hultborn, 2006). Since the 1960’s, PAD was thought to be activated by GABA_A_ receptors at the Ia afferent terminal, resulting in a shunting of terminal currents, depression of neurotransmitter release and ultimately a reduction in the excitatory postsynaptic potential (EPSP) evoked in the motoneuron, as reviewed in (Willis, 2006). However, the role of PAD and GABA_A_ receptors in producing presynaptic inhibition in the Ia afferent terminal has recently been disproven by multiple lines of evidence (Hari *et al*., 2021). For example, there is a sparsity of GABA_A_ receptors on the Ia afferent terminal (Alvarez *et al*., 1996; Hari *et al*., 2021) and correspondingly, PAD measured at the unmyelinated Ia afferent terminal is small-to-nonexistent and brief, lasting for 20 ms compared to more proximal, myelinated regions of the Ia afferent that contain nodes of Ranvier where PAD lasts for 100-200 ms (Lucas-Osma *et al*., 2018). Further, computer simulations confirm that PAD at the Ia afferent terminal is too weak and brief to produce a physiologically relevant decrease in action potential size to reduce EPSP activation in the motoneuron (Hari *et al*., 2021). In contrast, Ia afferent terminals are densely covered by GABA_B_ receptors that contribute to terminal presynaptic inhibition (Curtis & Lacey, 1994; Fink, 2013; Hari *et al*., 2021).

PAD and GABA_A_ receptors have instead been shown to facilitate sodium channels at the nodes of Ranvier in Ia afferents, helping spike propagation through branch points to facilitate motoneuron EPSPs (Hari *et al*., 2021; Metz *et al*., 2021). This is mediated by axoaxonic connections from GABAergic neurons (GABA_axo_ neurons) onto GABA_A_ receptors at or near nodes producing PAD that brings sodium spikes closers to threshold (Hari *et al*., 2021). Despite its minor influence at the Ia afferent terminal, GABA_A_ receptors have repeatedly been shown to contribute to the inhibition of the monosynaptic reflex in extensor motoneurons following conditioning of an antagonist flexor nerve to evoke PAD, since this inhibition is partly decreased by GABA_A_ receptor blockers (Eccles *et al*., 1963; Stuart & Redman, 1992; Curtis & Lacey, 1994; Curtis, 1998). This raises the question of how can the activation of GABA_A_ receptors be inhibitory in some instances (i.e., reduce reflexes) and excitatory in other (i.e., facilitate afferent nodal conduction), and how is this related to the observations that the extensor H-reflex is sometimes inhibited by an antagonist afferent conditioning stimulation (Mizuno *et al*., 1971; El-Tohamy & Sedgwick, 1983; Berardelli *et al*., 1987; Hultborn *et al*., 1987a; Nakashima *et al*., 1990; Burke *et al*., 1992; Capaday *et al*., 1995; Faist *et al*., 1996; Iles, 1996; Aymard *et al*., 2000; Howells *et al*., 2020) and at other times, the same H-reflex is facilitated by this conditioning stimulation (Hari *et al*., 2021; Metz *et al*., 2021)?

One intriguing possibility recently proposed is that PAD can induce post-activation depression in the Ia afferent terminals mediating the H-reflex (Hari *et al*., 2021). Here, we broadly define post-activation depression as a reduction in a monosynaptic EPSP caused by a prior activation of the same EPSP, where the first and second EPSPs are evoked by the same Ia afferent population each time, regardless of how the afferents are activated. Post-activation depression can be caused by numerous mechanisms, including transmitter depletion in the Ia afferents following the first EPSP, decreased afferent excitability from the post-spike refractory period that follows the first Ia afferent activation, and even inhibition of the afferent terminals by GABA_B_ receptors indirectly activated by the test afferents that activated the first EPSP (GABA_B_ mediated presynaptic inhibition) (Curtis & Eccles, 1960; Eccles *et* al., 1961a; Hultborn *et al*., 1996).

But how would PAD evoked from a conditioning stimulation to an antagonist afferent produce an EPSP in the agonist motoneurons and subsequent post-activation depression of EPSPs and related H-reflexes? Superficially this seems contrary to the conventional understanding that antagonist afferents generally produce reciprocal postsynaptic inhibition of agonist motoneurons (Sherrington, 1908). However, it is well known that PAD can produce antidromic action potentials in sensory axons that are recorded as dorsal root reflexes in the proximal afferent (termed dorsal root reflexes, DRRs), and PAD is strongest when evoked in extensor Ia afferents by antagonist flexor afferent conditioning (Eccles *et al*., 1961b; Rudomin & Schmidt, 1999). Further, since the early work of Eccles, it has been known that PAD in Ia afferents can also evoke *orthodromic* action potentials as evidenced by the generation of motoneuron EPSPs from these PAD-evoked spikes (Eccles *et al*., 1961b; Duchen, 1986; Willis, 1999). Theoretically, this orthodromic action potential evoked in the Ia afferent terminal should reduce subsequent neurotransmitter release for seconds via post-activation depression and thus, may result in a depression of subsequent EPSPs evoked by directly stimulating this same Ia afferent, just as the motoneuron EPSP is reduced by direct, repetitive activation of the Ia afferent. The latter EPSP depression with repetitive activation of the Ia afferent is termed rate dependent depression (RDD) and is caused by post-activation depression of the Ia afferent, and not terminal GABA_A_ receptor mediated presynaptic inhibition (Hultborn *et al*., 1996). This led us to speculate that H-reflexes following a conditioning stimulation of an antagonist nerve may be suppressed if the antagonist afferents activate PAD and orthodromic action potentials in the Ia afferents mediating the H-reflex, as suggested for the suppression of motoneuron EPSPs in the rodent when followed by PAD-evoked spikes (Duchen, 1986; Hari *et al*., 2021). To examine this, we first sought confirmation in mice that when orthodromic action potentials are activated in the Ia afferent from PAD circuits (PAD-evoked spikes), then subsequent direct activation of the Ia afferent produces a smaller EPSP in the motoneuron when tested within the time course of post-activation depression and related RDD (between 20 ms to 2s). For this we activated PAD either through an optogenetic activation of specific GABA neurons with axo-axonic connections to Ia afferents and their nodes (light activation of GAD2+ neurons in GAD2-cre//ChR2 mice), or from a separate, heteronymous sensory pathway (Hari et al., 2021). The former optogenetic method is especially powerful because any direct action of GABAergic neurons on motoneurons is inhibitory and so observations of EPSPs that are evoked by light unequivocally demonstrate that the PAD-evoked spikes drive these EPSPs.

We also examined in humans if a conditioning stimulation of the antagonist flexor nerve to the extensor soleus muscle, the common peroneal nerve (CPN), could likewise produce a short latency response in the soleus muscle indicative of activating an orthodromic, PAD-evoked spike in the soleus Ia afferent, as directly shown in sural and median afferents with microneurography (Shefner *et al*., 1992). We then examined if the suppression of subsequently activated H-reflexes was related to the presence or size of this putative PAD-evoked reflex at stimulation delays previously incorrectly attributed to the time-course of PAD-mediated presynaptic inhibition (30 to 400 ms). We also examined the time course of this antagonist-evoked H-reflex suppression at much longer delays within the post-activation depression window (500 ms to 2.5 s). The profile of the antagonist (flexor) H-reflex suppression was compared to the profile of RDD, and associated post activation depression, produced from repeated, direct activation of the soleus Ia afferents mediating the H-reflex at similar delays. We hypothesized that a similar long-lasting profile of H-reflex suppression from CPN conditioning or RDD would indicate that both were mediated by post-activation depression of the Ia afferents.

A more obvious, but rarely measured, mechanism of H-reflex suppression by antagonist conditioning stimulation is postsynaptic inhibition of the motoneuron given the well-known *glycinergic,* Ia-reciprocal postsynaptic inhibition on the motoneuron and the less well known inhibition caused by *GABA*, where 70-80% of GABAergic interneurons with projections to afferent terminals (GABA_axo_) also have projections onto the postsynaptic motoneuron (Pierce & Mendell, 1993; Hughes *et al*., 2005). Thus, we first examined if a brief conditioning stimulation of antagonist afferents directly inhibits the motoneuron with a time course similar to the long-lasting suppression of the H-reflex previously attributed to PAD. Inhibition of the soleus H-reflex from a brief vibration to the antagonist tibialis anterior (TA) tendon, or from percutaneous electrical stimulation to the common peroneal nerve (CPN), was measured at interstimulus intervals (ISIs) between 0 and 500 ms. Direct effects of the conditioning stimulation onto the test motoneuron(s) were measured from changes in the firing rate of tonically discharging single motor units in response to the conditioning stimulation alone. Changes in the firing rate of the tonically firing motor unit were taken as evidence of a postsynaptic effect on the motoneuron (Powers & Binder, 2001). We hypothesized that if the profile of suppression in the firing rate of the motor unit was similar to the profile of inhibition of the H-reflex from the same conditioning stimulation, then postsynaptic inhibition of the test motoneuron mediated a part of the H-reflex inhibition.

Parts of the data from the human vibration experiments (Hari *et al*., 2021) and the low-intensity CPN stimulation (Metz *et al*., 2021) have been published and are expanded upon here.

## Methods

### Participants, animals and ethics

Human experiments were approved by the Human Research Ethics Board at the University of Alberta (Protocol 00078057) and conducted with informed consent of the participants. Our sample comprised of 19 participants (7 male) with no known neurological injury or disease, ranging in age from 21 to 57 years (26.4 ± 10.2, mean ± standard deviation). *In vitro* recordings were made from adult mice (2.5-6 months old, both female and male equally) without (control) and with Cre expressed under the endogenous *Gad2* promotor region (*Gad2tm1(cre/ERT2)Zjh* mice abbreviated to Gad2-CreER mice, The Jackson Laboratory, Stock # 010702) as per Hari et al., 2021. These GAD2-CreER mice were crossed with mice containing flx-stop-flx-ChR2 expressed under the ubiquitous promotor Rosa, to yield mice with GAD2+ neurons expressing light sensitive ChR2 after they were injected with tamoxofin (abbreviated Gad2//ChR2 mice, as detailed in Hari et al. 2021). *In vivo* recordings were made from adult rats (3 - 8 months old, female only, Sprague-Dawley) as per Hari et al., 2021. All experimental procedures were approved by the University of Alberta Animal Care and Use Committee, Health Sciences division (Protocol AUP 00000224).

### Animal experimental setup

#### In vitro recordings in mice

Following extraction of the sacrocaudal spinal cord, dorsal and ventral roots (DR and VR) were mounted on silver-silver chloride wires above the nASCF of the recording chamber and covered with grease for monopolar stimulation and recording (see Hari et al., 2021 for details). Dorsal roots were stimulated with a constant current stimulator (Isoflex, Israel) with short pulses (0.1 ms) at 1.1 – 1.5 x threshold (T) to specifically activate proprioceptive afferents to evoke both PAD in Ia afferents and monosynaptic EPSPs in motoneurons. This grease gap method was also used to record the composite intracellular response of many sensory axons or motoneurons where the high impedance seal on the dorsal or ventral roots reduces extracellular currents, allowing the recording to reflect intracellular potentials (Luscher *et al*., 1979; Leppanen & Stys, 1997; Lucas-Osma *et al*., 2018). Return and ground wires were placed in the bath and likewise made of silver-silver chloride. Specifically for sensory axons, we recorded from the central ends of dorsal roots cut within about 2 - 4 mm of their entry into the spinal cord, to give the compound potential from all afferents in the root (dorsal roots potential, DRP), which has previously been shown to correspond to PAD (Lucas-Osma *et al*., 2018). The dorsal root recordings were amplified (2,000 times), high-pass filtered at 0.1 Hz to remove drift, low-pass filtered at 10 kHz, and sampled at 30 kHz (Axoscope 8; Axon Instruments /Molecular Devices, Burlingame, CA). These grease gap recordings of PAD on sensory afferents reflect only the response of the largest diameter axons in the dorsal root, mainly group I proprioceptive afferents, as detailed previously (Hari *et al*., 2021). The composite EPSPs in many motoneurons were likewise recorded from the central cut end of ventral roots mounted in the grease gap, which has also previously been shown to yield reliable estimates of the EPSPs (Fedirchuk *et al*., 1999). The EPSPs were identified as monosynaptic by their rapid onset (first component, ∼1 ms after afferent volley arrives in the ventral horn), lack of variability in latency (< 1 ms jitter), persistence at high rates (10 Hz) and appearance in isolation at the threshold for DR stimulation (< 1.1xT), unlike polysynaptic reflexes which vary in latency, disappear at high rates, and mostly need stronger DR stimulation to activate.

Light was used to evoke PAD in the GAD2//ChR2 mice and dorsal root stimulation was used to evoke PAD in control mice as described previously (Lin *et al*., 2019). Light was derived from a laser with a 447 nm wavelength (D442001FX lasers from Laserglow Technologies, Toronto) and was passed through a fibre optic cable (MFP_200/220/900-0.22_2m_FC-ZF1.25, Doric Lenses, Quebec City). A half cylindrical prism the length of about two spinal segments (8 mm; 3.9 mm focal length, Thor Labs, Newton, USA,) collimated the light into a narrow long beam (200 μm wide and 8 mm long). This narrow beam was focused longitudinally on the left side of the spinal cord roughly at the level of the dorsal horn, to target the epicentre of GABA_axo_ neurons, which are dorsally located. ChR2 rapidly depolarizes neurons (Zhang *et al*., 2011), and thus we used 5 - 10 ms light pulses to activate GABA_axo_ neurons, as confirmed by direct recordings from these neurons (Hari *et al*., 2021). Light was always kept at a minimal intensity, 1.1x T, where T is the threshold to evoke a light response in sensory axons, which made local heating from light unlikely.

Intracellular recordings of Ia afferent branches in the dorsal horn of rats were performed as in (Hari *et al*., 2021). Briefly, glass capillary tubes (1.5 mm and 0.86 mm outer and inner diameters, respectively; with filament; 603000 A-M Systems; Sequim, USA) with a bevelled hypodermic-shaped point of < 100 nm, were filled through their tips with 1 M K-acetate and 1 M KCl. Intracellular recording and current injection were performed with an Axoclamp2B amplifier (Axon Inst. and Molecular Devices, San Jose, USA). Recordings were low pass filtered at 10 kHz and sampled at 30 kHz (Clampex and Clampfit; Molecular Devices, San Jose, USA). Electrodes were advanced into myelinated afferents of the sacrocaudal spinal cord with a stepper motor (Model 2662, Kopf, USA, 10 μm steps at maximal speed, 4 mm/s), usually at the boundary between the dorsal columns and dorsal horn gray matter. Upon penetration, afferents were identified with direct orthodromic spikes evoked from DR stimulation. The lowest threshold proprioceptive group Ia afferents were identified by their direct response to DR stimulation, very low threshold (< 1.5 x T, T: afferent volley threshold), short latency (group Ia latency, coincident with onset of afferent volley), and antidromic response to micro stimulation of the afferent terminal in the ventral horn [∼ 10 μA stimulation via tungsten microelectrode as per (Lucas-Osma *et al*., 2018)]. Post hoc these were confirmed to be large proprioceptive Ia afferents by their unique extensive terminal branching around motoneurons, unlike large cutaneous Aβ afferents that do not project to the ventral horn. Clean axon penetrations without injury occurred abruptly with a sharp pop detected on speakers attached to the recorded signal, the membrane potential settling rapidly to near – 70 mV, and > 70 mV spikes usually readily evoked by DR stimulation or brief current injection pulses (1 – 3 nA, 20 ms, 1 Hz). Sensory axons also had a characteristic >100 ms long depolarization following stimulation of a dorsal root (primary afferent depolarization, PAD, at 4 - 5 ms latency) and short spike afterhyperpolarization (AHP ∼ 10 ms), which further distinguished them from other axons or neurons. Injured axons had higher resting potentials (> −60 mV), poor spikes (< 60 mV) and low resistance (to current pulse; Rm < 10 MΩ) and were discarded.

#### In vivo recordings in rats

The influence of evoking PAD in Ia afferents on the monosynaptic reflex (MSR) was performed in awake rats with percutaneous tail EMG recording and nerve stimulation as per Hari *et al*., 2021. Here, PAD was evoked by a cutaneous conditioning stimulation of the tip of the tail (0.2 ms pulses, 3xT, 40 - 120 ms prior to MSR testing) using an additional pair of fine Cooner wires implanted at the tip of the tail (separated by 8 mm). In rats the MSR latency is later than in mice due to the larger peripheral conduction time, ∼12 ms (as again confirmed by a similar latency to the F wave). This MSR was thus quantified by averaging the rectified EMG over a 12 - 20 ms window. Also, to confirm the GABA_A_ receptor involvement in regulating the MSR, the antagonist L655708 was injected systemically (1 mg/kg i.p., dissolved in 50 μl DMSO and diluted in 900 μl saline). Again, the MSR was tested at matched background EMG levels before and after conditioning (or L655708 application) to rule out changes in postsynaptic inhibition.

### Human experimental setup

Participants were seated in a reclined, supine position on a padded table. The right leg was bent slightly to access the popliteal fossa and padded supports were added to facilitate complete relaxation of all leg muscles because descending activation could potentially activate GABAergic circuits within the spinal cord (Jankowska *et al*., 1981; Rudomin, 1990; Eguibar *et al*., 1997; Ueno *et al*., 2018). During H-reflex recordings, participants were asked to rest completely with no talking, hand or arm movements.

#### Surface EMG recordings

A pair of Ag-AgCl electrodes (Kendall; Chicopee, MA, USA, 3.2 cm by 2.2 cm) was used to record surface EMG from the soleus and tibialis anterior (TA) muscles with a ground electrode placed just below the knee. The EMG signals were amplified by 1000 and band-pass filtered from 10 to 1000 Hz (Octopus, Bortec Technologies; Calgary, AB, Canada) and then digitized at a rate of 5000 Hz using Axoscope 10 hardware and software (Digidata 1400 Series, Axon Instruments, Union City, CA). To examine the postsynaptic effects of the conditioning inputs alone on the soleus motoneurons, surface EMG electrodes were also used to record single motor units during weak contractions by placing the surface electrodes on the lateral border of the muscle as done previously (Matthews, 1996). Single motor units were also recorded using a high-density surface EMG electrode (HDsEMG, OT Bioelettronica, Torino, Italy, Semi-disposable adhesive matrix, 64 electrodes, 5×13, 8 mm inter-electrode distance) with ground and differential electrodes wrapped around the ankle and below the knee. Signals were amplified (150 times), filtered (10 to 900 Hz) and digitized (16 bit at 5120 Hz) using the Quattrocento Bioelectrical signal amplifier and OTBioLab+ v.1.2.3.0 software (OT Bioelettronica, Torino, Italy). The EMG signal was decomposed into single motor units using custom MatLab software as per (Negro *et al*., 2016; Afsharipour *et al*., 2020).

#### Nerve stimulation to evoke an H-reflex

The tibial nerve (TN) was stimulated in a bipolar arrangement using a constant current stimulator (1 ms rectangular pulse width, Digitimer DS7A, Hertfordshire, UK) to evoke an H-reflex in the soleus muscle. After searching for the TN with a probe to evoke a pure plantarflexion, an Ag-AgCl electrode (cathode: Kendall; Chicopee, MA, USA, 2.2 cm by 2.2 cm) was placed in the popliteal fossa, with the anode (Axelgaard; Fallbrook, CA, USA, 5 cm by 10 cm) placed on the patella. Stimulation intensity was set to evoke a test H-reflex of approximately half maximum on the ascending phase of the H-reflex recruitment curve to allow for both facilitatory and inhibitory effects of the conditioning input to be revealed (Crone *et al*., 1990). H-reflexes recorded at rest were evoked every 5 seconds to minimize post-activation depression from RDD (Hultborn *et al*., 1996) and at least 10 H-reflexes were evoked before conditioning to establish a steady baseline. All H-reflexes were recorded at rest.

#### Antagonist TA tendon vibration

In 8 participants, the soleus H-reflex was conditioned by a prior vibration of the TA tendon (3 pulses, 200 Hz) to preferentially activate Ia afferents as done previously (Hultborn *et al*., 1987a). A 7 mm diameter probe attached to an audio amplifier was pressed gently against the TA tendon at the base of the leg. Approximately 10 baseline soleus H-reflexes were elicited to ensure the H-reflex was stable, followed by 7 conditioned H-reflexes at a single ISI (0, 30, 60, 100, 150, 200, 300, 400 or 500 ms). Following this, 3 to 4 unconditioned H-reflexes were evoked to reestablish baseline and another run of 7 conditioned H-reflexes was applied at a randomly chosen interval. This was repeated until all ISI intervals were applied (Fig. 1A).

**Figure 1:**
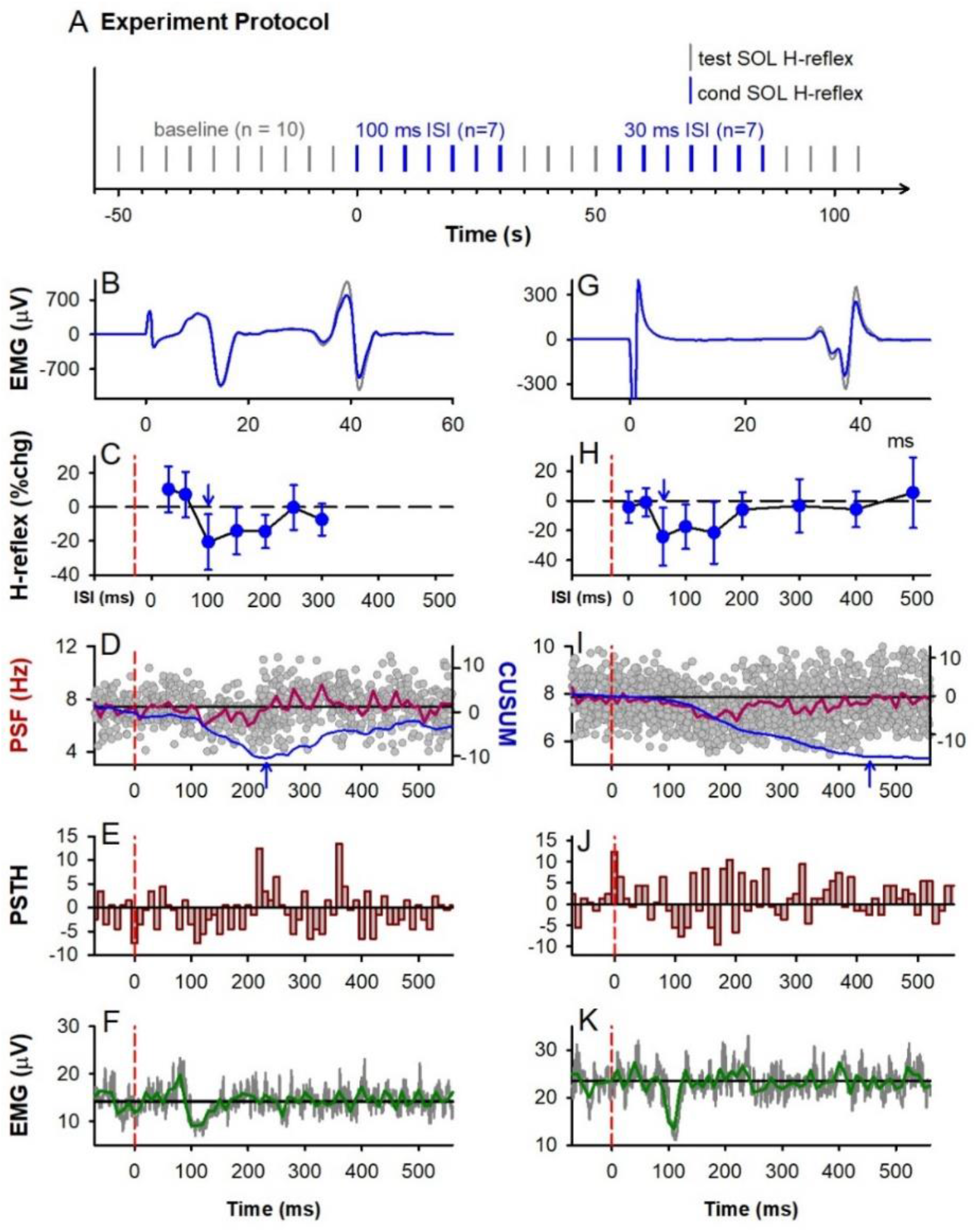
TA tendon vibration. **A)** General experiment schematic: vertical bars mark application of unconditioned test (grey) and conditioned (blue) H-reflexes to the soleus (SOL) muscle, evoked every 5 s with random application of the various conditioning ISIs interposed with a run of test H-reflexes. **B-F** & **G-K**: Representative data from 2 participants. **B,G)** Average of 7 test (grey) and 7 conditioned (blue) soleus H-reflexes (unrectified EMG) at the 100 ms (B) and 60 ms (G) ISI respectively. TN stimulation to evoke H-reflex applied at 0 ms. **C,H)** % change (chg) of the soleus H-reflex (mean ± SD) plotted at each ISI with peak % change indicated by the arrow. Time of vibration indicated by red dashed line, occurring to the left of the 0 ms ISI by the latency of H-reflex. **D,I)** Superimposed firing rates (grey dots, PSF) of soleus motor units in response to TA tendon vibration alone (185 and 340 sweeps respectively), with a 10-ms binned average rate (red line, mean PSF) and mean PSF CUSUM (blue line). **E,J)** Post-stimulus time histogram (PSTH) showing motor unit count per each 10 ms bin with mean pre-vibration count subtracted. **F,K)** Rectified soleus EMG (grey lines, 31 and 68 sweeps respectively), with 10-ms binned average (green line). Vertical dashed red lines in C to K mark onset of TA tendon vibration (at time 0). Horizontal black lines mark the average pre-vibration values (from −100 to 0 ms).

Modulation in the tonic firing rate of single soleus motor units was used to determine if the TA tendon vibration had any postsynaptic actions on the soleus motoneurons. A small voluntary isometric contraction (∼5% of maximum) was used to produce a steady discharge of the single unit(s) using both auditory and visual feedback, while the vibration was delivered every 3 seconds. Seven units from 5 participants were isolated from the surface EMG and 12 units from 3 participants were decomposed from the HDsEMG. Modulation of the whole-muscle surface EMG was also measured while participants held a stronger isometric voluntary contraction at approximately 10% of maximum.

#### Antagonist CPN stimulation

In a separate experiment on a different day in 13 participants (2 participants from the vibration experiment), the CPN supplying the antagonist TA muscle was stimulated to condition the ipsilateral soleus H-reflex as done previously (Mizuno *et al*., 1971; El-Tohamy & Sedgwick, 1983; Capaday *et al*., 1995). The CPN was stimulated using a bipolar arrangement (Ag-AgCl electrodes, Kendall; Chicopee, MA, USA, 2.2 cm by 2.2 cm) with the anode placed anterior and slightly distal to the fibular head on the right leg and the cathode 2-3 cm more proximal. Care was taken to elicit a pure dorsiflexion response without foot eversion. Three pulses (200 Hz, 1-ms pulse width) were applied to the CPN at an intensity of 1.0 and 1.5 x motor threshold (MT) at the 3, 15, 30, 60, 80, 100, 150, 200, 300 and 400 ms ISIs using a similar protocol as in the vibration experiment (Fig. 1A). Motor threshold was determined by the lowest-intensity, single-pulse stimulation of the CPN that produced a small (< 10 μV) but reproducible direct motor response (M wave) in the TA muscle at rest. In a separate trial, CPN stimulation was applied at longer ISI intervals (500, 1000, 1500, 2000 and 2500 ms) at both the 1.0 and 1.5 x MT stimulation intensities with the same protocol as for the shorter ISIs.

The effect of the 1.0 and 1.5 x MT conditioning CPN stimuli on the firing rate of tonically discharging soleus motor units and whole muscle EMG were also measured as done for the vibration stimuli described above. In 12 participants, 44 units were isolated from the surface EMG and in 1 participant, 2 units were decomposed from the HDsEMG.

#### Post-activation depression of the soleus H-reflex during RDD

In 8 of the 13 participants from the CPN-conditioning experiment, we examined the suppression of the soleus H-reflex in response to repetitive stimulation of the TN to directly assess post-activation depression and compare it to the long-lasting depression evoked from a conditioning heteronymous CPN stimulation. To measure post-activation depression during RDD, the first H-reflex (H1) of a stimulation trial was evoked at approximately 50% of maximum on the ascending part of the H-reflex recruitment curve. A run of at least 10 H-reflexes were repetitively evoked at the same conditioning-test intervals as for the CPN stimulation at 500, 1000, 1500, 2000 and 2500 ms. Three trials were performed for each stimulation frequency with at least 30 to 40 s in between each trial.

### Data analysis

#### EPSP modulation

Similar to Hari et al., 2021, the amplitude of the EPSP recorded in the ventral root evoked from dorsal root stimulation was compared with and without action potentials evoked in the Ia afferent from optogenetic or sensory activated PAD. The average EPSP from ∼10 trials evoked every 10 s just before conditioning and 10 trials during conditioning was used to compute the change in the peak size of the monosynaptic EPSP with conditioning. The background motoneuron potential, membrane resistance (Rm) and time constant just prior to the EPSP was also assessed before and after conditioning to examine whether there were any postsynaptic changes that might contribute to changes in the EPSP with conditioning. The latency of the direct EPSPs evoked by dorsal root stimulation was also compared to the latency of the EPSPs from PAD-evoked spikes. Along with the ventral root recordings, PAD was simultaneously recorded from the dorsal roots by a similar averaging method (10 trials of conditioning), to establish the relation of changes in EPSPs with associated PAD.

#### H-reflex modulation

For a given trial run, the average, unrectified peak-peak amplitude of all test (unconditioned) soleus H-reflexes was compared to the average peak-peak amplitude of the 7 conditioned soleus H-reflexes (vibration and CPN stimulation) for each of the ISIs tested. The conditioned H-reflexes were expressed as a % change from the test reflex using the formula: % change H-reflex = ([(condition H - test H)/test H]*100%). The mean % change of the soleus H-reflex at each ISI was then averaged across participants.

#### Effect of conditioning stimulation on soleus motoneurons

The firing rate profile of soleus motor units in response to the conditioning TA tendon vibration or CPN stimulation were superimposed and time-locked to the time of the stimulation (set to 0 ms) to produce a peri-stimulus frequencygram (PSF), as done previously (Norton *et al*., 2008). The PSF was divided into 10 ms bins and the mean of each bin was expressed as a percentage of the mean, pre-stimulus firing rate measured from a 100 ms window before the conditioning stimulation using the formula: % change PSF = ([(post-stimulus rate - pre-stimulus rate)/pre-stimulus rate]*100%). The mean % change in each bin was then averaged across participants to produce the group mean PSF. The cumulative sum (CUSUM) of the mean PSF was measured by subtracting the mean pre-stimulus values and integrating over time. A significant increase or decrease in the mean PSF CUSUM was considered when it fell above or below, respectively, 2 standard deviations (SD) of the mean pre-stimulus value. For group changes in EMG activity, a similar bin analysis was done for the rectified surface EMG. The mean pre-stimulus EMG measured between - 300 to 0 ms was subtracted from each of the mean bin values to measure the net increase or decrease in EMG activity. The artifact from the CPN stimulation (typically from 0 to 20 ms) was manually removed from the soleus EMG.

#### Post-activation depression of H-reflexes

To quantify the amount of post-activation depression during the RDD trials, the average peak-peak amplitude of the second to eighth H-reflex (H2-H8) was compared to the peak-peak amplitude of the first H-reflex (H1) in each trial run using the formula: % change post-activation depression = ([(H2-H8)/H1]*100%). The % change post-activation depression value for the three trials at each stimulation frequency was averaged together and this value was then averaged across the 8 participants. The resulting % change in post-activation depression at each stimulation frequency was compared to the % change of the CPN-conditioned soleus H-reflex at the corresponding ISI.

### Statistical Analysis

All statistics were performed with SigmaPlot 11 software. To determine if the suppression of the H-reflex across the various conditioning stimulation intervals was different from a 0% change, a One-Way Repeated Measure ANOVA was used because the data was normally distributed. The F value was reported and post hoc Tukey Tests were used to determine which ISIs were significantly different from a 0% change. A Two-Way Repeated Measure ANOVA was used to compare if the H-reflex suppression using antagonist CPN stimulation was different across the various ISIs compared to the repetitive TN conditioning stimulation during RDD. Student t-tests and Pearson Product-Moment Correlation were used to compare the two groups of normally distributed data. Data are presented in the figures and in the text as means ± standard deviation (SD). Significance was set as P < 0.05.

## Results

### Antagonist TA tendon vibration

We started by examining the action of tendon vibration, as this was the classic method of examining presynaptic inhibition of Ia afferents in humans (Morin *et al*., 1984; Hultborn *et al*., 1987a; Hultborn *et al*., 1987b; Roby-Brami & Bussel, 1990; Rossi *et al*., 1999). The suppression of the soleus H-reflex from a prior vibration (3 cycles at 200 Hz) to the antagonist TA tendon was examined in 8 participants (schematic of experimental protocol in Fig. 1A). Similar to earlier studies, the soleus H-reflex was suppressed between 60 to 400 ms following the brief tendon vibration, as shown for participant 1 (Fig. 1B-C) and participant 2 (Fig. 1G-H). Across the 8 participants, the amplitude of the conditioned soleus H-reflex was modulated across the various ISIs, with the H-reflex being significantly suppressed at the 100, 150 and 200 ms ISIs (marked by white circles, Fig. 2A, see statistics in legend).

**Figure 2:**
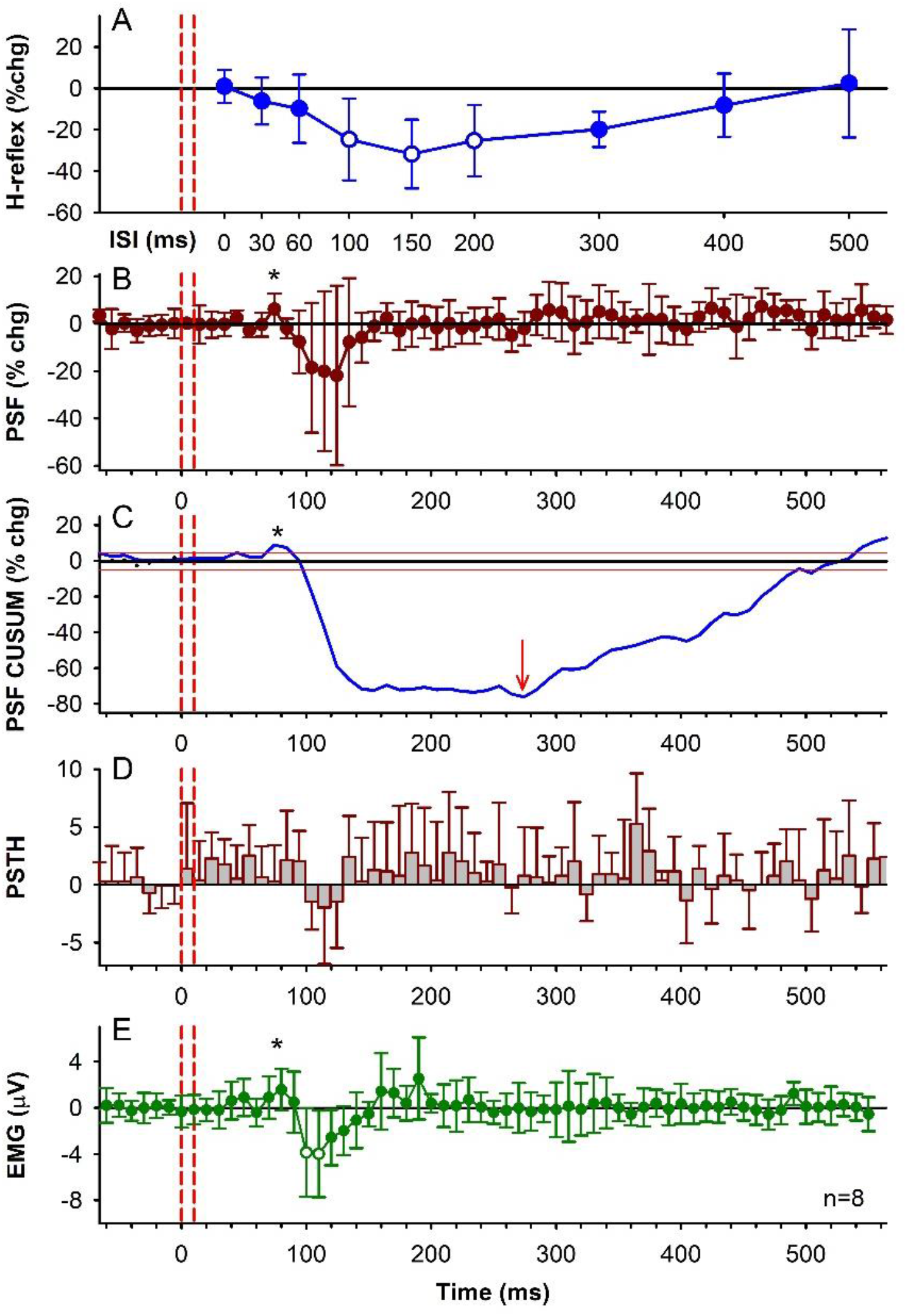
TA tendon vibration, group data. **A)** % change (chg) of the soleus H-reflex from TA tendon vibration at each ISI averaged across the group (n=8 participants). The average size of the unconditioned test soleus H-reflex was 1581 ± 648 μV (50.3 ± 15.5% of H-Max). There was an effect of ISI on the conditioned H-reflexes [F(7, 9)=45.2, P<0.001, one-way RM ANOVA] with post-hoc analysis showing the 100 ms (P = 0.015), 150 ms (P = 0.005) and 200 ms (P = 0.012) ISIs significantly different from a 0% change (white symbols). The 0 ms ISI is shifted to the right of the vibration onset (first dashed red line) by the average latency of the H-reflex. **B)** % change of the PSF averaged across the group (10-ms bins) in response to TA tendon vibration alone. There was an effect of time on the % change PSF (F[71,64] = 2.02, *P* < .001, one-way RM ANOVA), but no single value was significantly different than a 0% change (post hoc Tukey Test, P >0.05). **C)** CUSUM of mean PSF in B, plotted with the mean pre-vibration value (horizontal black line) and 2 SD values above and below the mean line (red lines). **D)** Number of soleus motor unit action potentials averaged across the group in each 10-ms bin (PSTH) with the mean pre-vibration number subtracted. The count per bin did not significantly change across the time bins (F[71,64] = 1.10, *P* = 0.288, one-way RM ANOVA). **E)** Rectified soleus EMG (mean EMG - pre-stimulus EMG) in each 10-ms bin averaged across the group. The EMG was significantly different across the tested time bins (F[7,56] = 2.85, *P* < 0.001, one-way RM ANOVA), being significantly different from 0 mV at the 100 ms (P = 0.015) and 110 ms (P = 0.009, Tukey Test, white symbols) bins. Vertical red dashed lines indicate the onset and offset of TA tendon vibration. Error bars indicate ± SD. * indicates small increases in motor unit or EMG activity shortly after the vibration.

To determine if the profile of H-reflex suppression was mediated, in part, from a postsynaptic inhibition of the soleus motoneurons, the firing rate of a voluntarily activated soleus motor unit was measured in response to the conditioning tendon vibration applied alone. Following the conditioning stimulus, the mean firing rate of the soleus motor unit measured from the peristimulus frequencygram (PSF) decreased below the mean pre-stimulus rate, with a time course similar to the soleus H-reflex inhibition (between 80 and 400 ms in Figs. 1D & I, red trace). Note that the H-reflex data plotted at the various ISIs are shifted to the right of the PSF by ∼ 30 ms, the latency of the H-reflex, to estimate the excitability of the spinal motoneurons at the time they were activated by Ia afferents in the H-reflex (see also Metz *et al*., 2021). The decrease in the PSF is an indication of a prolonged inhibitory postsynaptic potential (IPSP) (Turker & Powers, 1999), with the duration of the IPSP marked by the lowest point in the blue PSF CUSUM line (Figs 1D and I). On average, the PSF was suppressed out to 270 ms (red arrow in PSF CUSUM, Fig. 2C) at time points when the H-reflex was also suppressed. There was also a decrease in the number of motor unit action potentials compared to the mean pre-stimulus count during the early decrease in the PSF, as assessed from the PSTH (Figs. 1E&J, Fig. 2D), that was also reflected in the decreased surface EMG during this period (Figs. 1 F&K, 2E). In some cases, the IPSP was large enough to produce a synchronous resetting of the tonic motor unit discharge, as marked by repetitive clusters of increased firing in the PSTH following the reduced firing periods (near 230 and 360 ms in Fig. 1E). Taken together, our results provide evidence for prolonged inhibition of the soleus motoneurons from the conditioning TA tendon vibration, which likely contributed to the profile of H-reflex suppression.

Unexpectedly, in some participants there was a clear early excitation of the extensor soleus EMG (abbreviated *early soleus reflex*) following vibration to the *antagonist TA* tendon applied alone (e.g., Fig. 1F). This excitation was evident in the group EMG averages and in the PSF and PSF CUSUM (at *, Figs. 2B,C and E), occurring at about 50 - 70 ms after the tendon vibration. The relevance of this early soleus excitation evoked by the antagonist afferent conditioning in suppressing the H-reflex was investigated further below.

### Post-activation depression of afferent transmission in mice and rats

The early soleus reflex following the TA tendon vibration raised the possibility that the antagonist TA afferents evoked a PAD in the soleus Ia afferents that produced orthodromic action potentials (PAD-evoked spikes) to subsequently activate a monosynaptic reflex (MSR), causing post-activation depression of soleus Ia afferent transmission, as we detail in the Introduction. Thus, we examined if GABAergic neurons with axo-axonic connections to afferents (GABA_axo_ neurons) could, in principle, evoke PAD and orthodromic spikes in Ia afferents to produce such post-activation depression and a reduction of subsequent motoneuron EPSPs. To do this, grease gap recordings from sacral dorsal and ventral roots were made in adult spinal cords from GAD2//ChR2 positive mice (schematic in Fig. 3A). These recordings allowed simultaneous assessment of afferent PAD and motoneuron EPSPs, with PAD directly evoked by GABA_axo_ (GAD2^+^) neurons, which cannot themselves directly evoke EPSPs in motoneurons (Hari *et al*., 2021). When light activation of GABA_axo_ neurons produced a PAD without generating action potentials in the afferents (top light blue trace, Fig. 3A), the monosynaptic EPSP evoked from a dorsal root (DR 1) stimulation during this PAD was facilitated (bottom light blue trace) compared to when PAD was not present (pink trace, DR 1 stim alone). As we have recently shown, this EPSP facilitation is due to increased conduction in Ia afferents evoking the EPSP, via PAD facilitating spike generation at the nodes [nodal facilitation (Hari *et al*., 2021; Metz *et al*., 2021)]. However, when activation of the GABA_axo_ neurons produced a larger and faster rising PAD that also generated an action potential in the Ia afferent (PAD-evoked spike, DRR; top black trace Fig. 3A), a fast transient EPSP was evoked in the motoneurons (middle black trace, Fig. 3A), indicating that the PAD-evoked spike traveled orthodromically to the Ia afferent terminal and evoked a monosynaptic EPSP, as Duchen (1986) has previously demonstrated. Indeed, this PAD-evoked spike in the Ia axon activates the motoneuron synchronously at a monosynaptic latency that is similar to the latency of EPSPs generated by direct activation of the Ia afferent by dorsal root stimulation (DR 1, Fig. 3F,G). Following these PAD-evoked spikes and EPSPs, subsequently tested monosynaptic EPSPs evoked by direct DR stimulation were always depressed, regardless of whether the EPSP was evoked during or after PAD and the related PAD-evoked EPSPs (Figs. 3A and B respectively, black traces), demonstrating that PAD-evoked spikes cause post-activation depression of the Ia-mediated EPSP (see also Fig. 3C).

**Figure 3:**
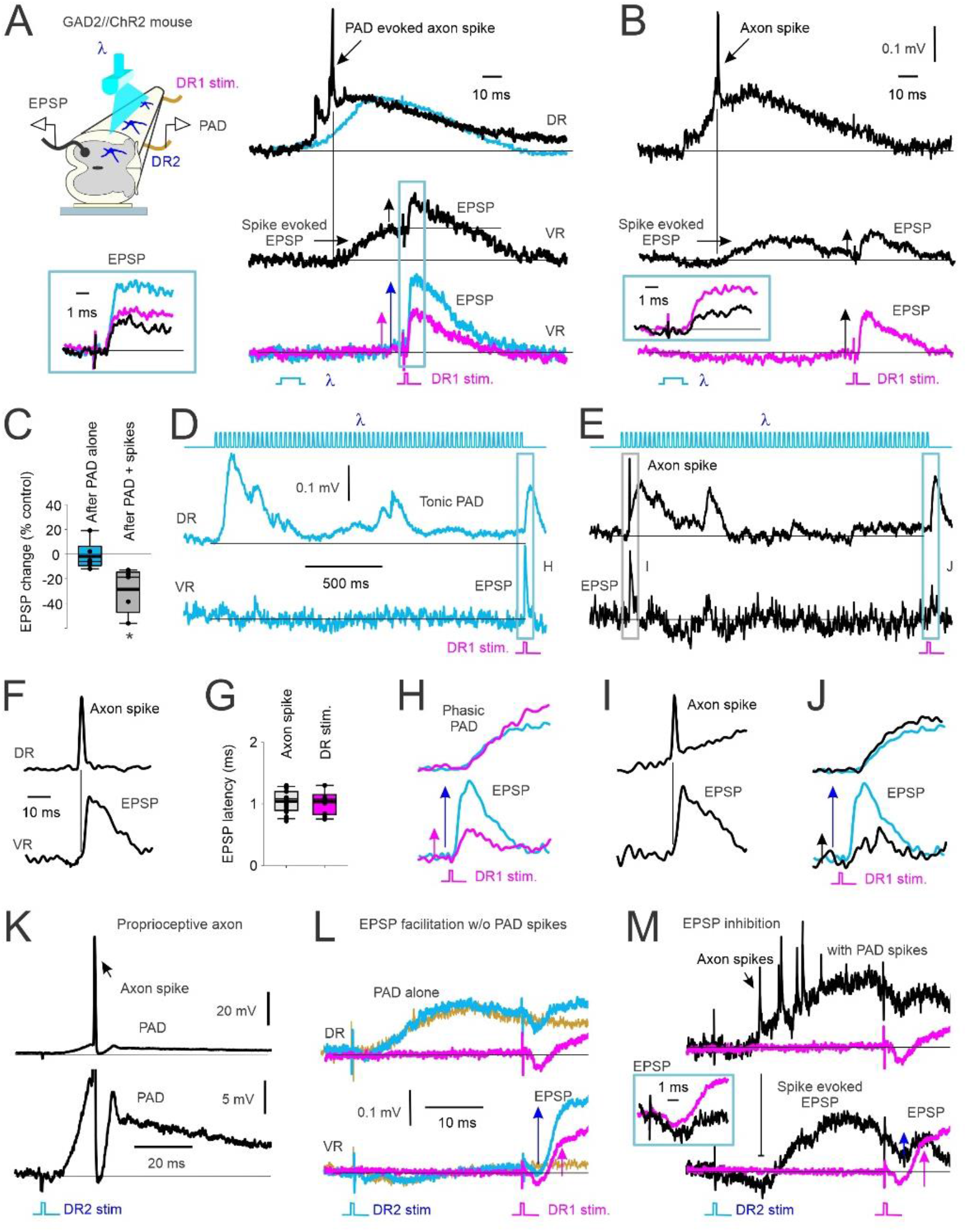
GABA_A_ mediated PAD in Ia afferents with and without spiking and subsequent EPSPs. **A)** *Left*, In vitro experimental set up showing PAD recorded from dorsal root 2 (DR2) activated by light (λ) in a GAD2//ChR2 positive mouse. Monosynaptic reflex (MSR) activated by stimulation of DR1, subsequent EPSPs recorded from ventral root (VR). *Right, top*: PAD recorded from DR2 with (black) and without (blue) PAD-evoked axon spike. *Middle*: VR recording of evoked EPSP from the PAD-evoked axon spike in top trace. Subsequent test EPSP activation (DR1 stimulation) is reduced (vertical arrows indicate EPSP amplitude). *Bottom*: EPSP from DR1 stimulation alone (pink) compared to EPSP following the light-evoked, non-spiking PAD shown in top trace (blue). *Box*: Overlay of EPSPs from middle and bottom traces in A. **B)** Same as in A but test EPSP from DR1 stimulation delivered at a longer interval after the PAD spike-evoked EPSP returned to baseline. *Box:* Overlay of conditioned (black) and control (pink, DR1 stimulation only) EPSPs. **C)** Change in amplitude of light-conditioned EPSPs (100 - 150 ms ISI) after a non-spiking PAD (PAD alone) and after a PAD with spike(s) as a percentage of an EPSP evoked from DR1 stimulation alone (control). Box plots: median, thin line; mean, thick line; interquartile range IQR, box bounds; most extreme data points within 1.5 x IQR, standard error bars. * = significantly different than 0% change, Mann Whitney U test, p < 0.05, n=5 mice. **D-E)** *Top*: Light-evoked tonic PAD recorded from DR2 (middle trace) without (D, blue) and with (E, black) a PAD-evoked spike. *Bottom*: VR recordings of associated of monosynaptic EPSP from DR1 stimulation and PAD spike-evoked EPSP. **F)** Example DR and VR recording of PAD-evoked axon spike and resulting EPSP at a monosynaptic latency on expanded time scale. **G)** Comparison of EPSP latency following PAD evoked axon spike and DR 1 stimulation. **H-J)** Expanded view of PAD (top) and EPSPs (bottom) evoked in D and E. Similar results observed in in n=6/6 mice. **K-M)** Same as in A and B but DR2 stimulation (1.5 x T) used to evoke PAD instead of light. Similar results in n = 5/5 rats. **K)** *Top*: Rat intracellular (IC) recording of proprioceptive (Ia) afferent with PAD-evoked axon spike from DR2 stimulation. *Bottom*: expanded vertical axis of top trace. **L-M)** PAD evoked with DR2 stimulation. *Top*: DR recording. *Bottom:* VR recording. **L)** PAD evoked without axon spikes. Pink: EPSP alone from DR1 stim. Blue: Non-spiking PAD evoked from DR2 stimulation and facilitated EPSP from DR1 stimulation. Yellow: PAD from DR2 stimulation applied alone. **M)** PAD, axon spikes and motoneuron EPSPs evoked from DR2 stimulation and test EPSP from DR1 stimulation without (pink) and with (black) spiking PAD (EPSPs expanded in box).

If the suppression of the monosynaptic EPSP following the PAD-evoked spike is indeed due to post-activation depression of the Ia afferent terminal, this suppression should last for many seconds (Curtis & Eccles, 1960; Hultborn *et al*., 1996). In support of this, we evaluated a very long interval between the PAD-evoked spikes and the test EPSP. When an EPSP was evoked during a tonic PAD produced by a long train of light pulses, the EPSP was facilitated by the PAD when no PAD-evoked spikes were produced, as previously reported (Hari et al., 2021) (Fig. 3D and expanded in H, pink trace EPSP alone, blue trace EPSP with tonic PAD). In contrast, when the same long train of light pulses evoked a spike at the start of the tonic PAD (Fig. 3E and expanded in I), the test EPSP that was evoked 1.5 s later was instead depressed (Fig. 3E and expanded in J, black trace), especially compared to the expected facilitation at this time (blue trace overlayed from Fig. 3H). This is broadly similar to Fig. 3 of Duchen 1986, though in that case PAD was evoked indirectly by a DR stimulation, as we detail next.

A similar suppression of the monosynaptic EPSP occurred when PAD-evoked spikes in the Ia afferent were activated by sensory-evoked PAD, rather than light-evoked PAD, the former produced by stimulating a separate dorsal root (termed DR2) to that used to evoke the test EPSP (DR1). As shown from an intracellular (IC) axon recording of a proprioceptive Ia afferent (top, Fig. 3K), a single dorsal root pulse (DR2) sometimes produced a spike in the Ia afferent on the rising phase of the PAD evoked by this dorsal root stimulation (expanded view in bottom). The same DR2 stimulation produced multiple axon spikes in the afferent population recorded from the dorsal root (top, Fig. 3M) that in turn evoked multiple compounded motoneuron EPSPs recorded in the ventral root (VR, bottom), as with ontogenetically evoked PAD (Fig 3A). Following these PAD-evoked EPSPs, a direct test EPSP evoked by a DR1 stimulation was suppressed (inset in Fig. 3M), even though it was evoked when the membrane potential of the motoneuron returned to near baseline. Similar to light activation of GABA_axo_ neurons when PAD did not produce axon spikes or associated depolarizations (EPSPs) of the motoneurons (DR2 stimulation only: yellow trace, Fig. 3L), then the test EPSP was facilitated by PAD (EPSP evoked by a DR 1 stimulation 1.5xT; blue trace, Fig. 3L) compared to the EPSP tested alone without PAD (pink trace, Fig. 3L). This absence or presence of PAD-evoked spikes occurred randomly at a fixed DR2 stimulation intensity, as in Figures 3L or M, respectively, though these spikes could also be reduced in occurrence by reducing the stimulation intensity and again, the EPSP was inhibited only when they occurred (not shown). In summary, the same sensory conditioning stimulation can produce either suppression or facilitation of subsequent EPSPs depending on whether the associated PAD activates orthodromic spikes in the Ia afferent. Additionally, afferents from one nerve (e.g., DR 2) can produce widespread activation of PAD in afferents from other nerves (DR 1) (Lucas-Osma *et al*., 2018), reminiscent of reflex irradiation (Zehr & Stein, 1999).

To explore whether PAD-evoked spikes can produce motoneuron reflexes *in vivo*, we first examined the special characteristics of these spikes and the reflexes they evoke *in vitro* so they can be distinguished from polysynaptic reflexes activated from other sources and ultimately be detected *in vivo.* We started by determining the origin of the PAD-evoked spikes by recording intracellularly from single group Ia afferents in the spinal cord near the dorsal root entry zone (Fig. 4A). A brief DR stimulation that was subthreshold to the Ia afferent being recorded (as indicated by a lack of an orthodromic spike at the green arrowhead in Fig. 4B) evoked a PAD in the recorded Ia afferent. In some of the trials, the DR stimulation produced one or more spikes starting on the rising phase of the PAD (dark blue Fig. 4B), which propagated antidromically toward the recording electrode and out the dorsal root and was recorded as a dorsal root reflex (DRR; Lucas-Osma et al., 2018). In trials where the DR stimulation did not produce these antidromic spikes, there were underlying oscillations on top of the PAD at the time where the spikes were in other trials (light blue, Fig. 4B). Previously, we have demonstrated that these oscillations represent spikes that are generated by PAD deep in the spinal cord (distal to the recording electrode) that fail to propagate antidromically out the dorsal root, but likely propagate orthodromically toward the motoneurons (Lucas-Osma *et al*., 2018), and so have functional importance. One key characteristic of these PAD-evoked spikes and oscillations is that they are blocked by hyperpolarization, with a small hyperpolarization of the Ia afferent consistently eliminating the spikes (pink, Fig 4B) and a stronger hyperpolarization eliminating the underlying oscillations (Fig. 4D).

**Figure 4.**
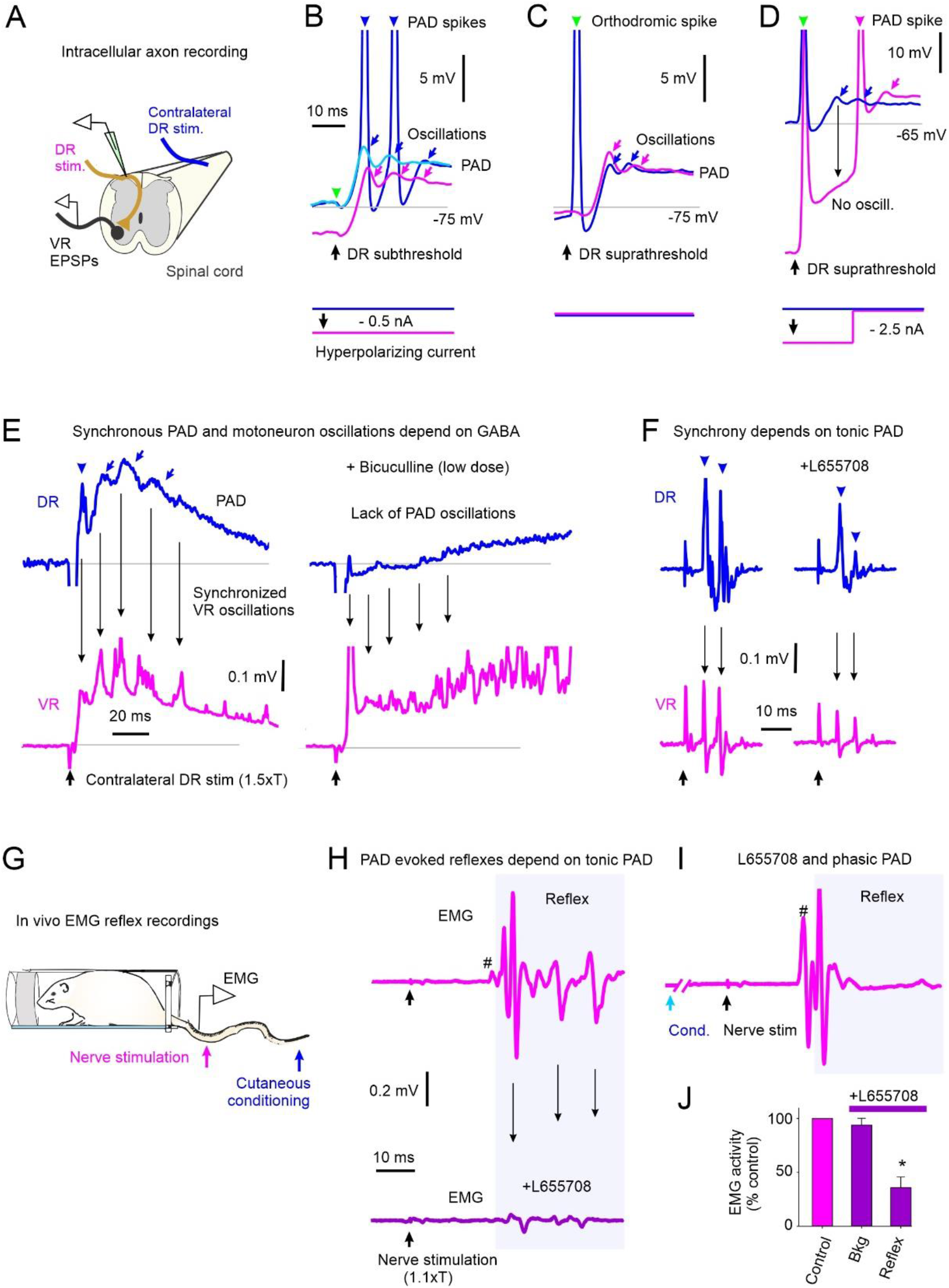
GABA_A_ mediated synchronous PAD and motoneuron oscillations. **A-D)** PAD recorded intra-axonally from a rat Ia afferent. **A)** In vitro experimental set up. **B)** Stimulation of DR that is subthreshold to orthodromic spike for recorded Ia afferent without (blue & light blue traces) and with (pink) a small, applied hyperpolarization. PAD-evoked spikes and oscillations marked by arrowheads and arrows, respectively. Predicted timing of non-activated orthodromic spike indicated by green arrowhead. **C)** Suprathreshold DR stimulation producing orthodromic spike (green arrow and blue trace) compared to PAD oscillations produced with subthreshold DR stimulation in A (pink). **D)** Suprathreshold DR stimulation with strong hyperpolarization of Ia afferent to reduce PAD oscillations from distal spikes, n = 10/10 axons similar to results in B-D. **E-F)** *Left:* Synchronized PAD (DR recording, top) and motoneuron EPSP (VR recording, bottom) oscillations from PAD evoked by contralateral DR stimulation in rat. *Right:* Application of the wide spectrum GABA_A_ receptor antagonist bicuculine **(E)** or the a5 GABA_A_ receptor antagonist L655708 **(F)** to the recording bath, similar findings in 7/7 rats. **G)** PAD evoked by stimulating cutaneous afferents in tip of tail in awake rat and MSR EMG recordings from more proximal tail nerve stimulation. **H)** *Top*: Small MSR (#) and oscillating EMG reflex responses from tail nerve stimulation (1.1xT) to recruit proprioceptive afferents. *Bottom*: EMG response to tail nerve stimulation following i.p. application of L655708. **I)** Facilitation of MSR and later reflex from prior (60 ms ISI) conditioning cutaneous stimulation of tip of tail. **J)** Pre-stimulus background (Bkd) EMG and post-MSR reflex activity (shaded window in I) following application of L655708 as a percentage of pre-drug (control). * = significantly different from control (Mann Whitney U test, p < 0.001, n = 6 rats).

Consistent with the recorded oscillations being mediated by distal sodium spikes, we found that inducing sodium spike inactivation by evoking a direct orthodromic spike in the Ia afferent via a supra-threshold stimulation to its DR (at green arrowhead, Fig. 4C) reduced the subsequent PAD-triggered oscillations measured in the DR (dark blue, Fig. 4C). Furthermore, after blocking these oscillations with a strong hyperpolarization, they resumed when the hyperpolarization was removed (Fig. 4D), the latter showing that they are mediated intrinsically to the axon and not by synaptic inputs. Another characteristic of the PAD-evoked spikes and oscillations is their unique timing, starting at about a 5-7 ms latency on the rising portion of a phasic PAD evoked by sensory stimulation (Fig. 4E, top left), and sometimes repeating at about 7-10 ms intervals, driving a characteristic oscillation in the population recording of afferents in the DRs, with a frequency (100 - 140 Hz) that is likely determined by the axon’s refractory period and associated AHP (seen in Fig. 4B). Finally, as with direct hyperpolarization of the Ia afferent, indirect afferent hyperpolarization induced by blocking spontaneous tonic PAD on the afferents with either a low dose of the non-selective GABA_A_ receptor antagonist bicuculline (Fig. 4E, top traces) or the selective extra-synaptic α5 GABA_A_ receptor antagonist L655708 (Fig. 4F, top traces; see Lucas-Osma et al 2018 for details) reduced PAD-evoked spikes and oscillations.

These same activation characteristics appear in the motoneurons (VR recordings, pink) when simultaneously recorded with the PAD, consistent with these PAD-evoked spikes driving the motoneuron EPSPs. That is, the population of motoneuron EPSPs in the VR recording followed the oscillatory profile of the PAD measured on the DR recording with the same timing and were eliminated by hyperpolarizing the afferents with bicuculline (Fig. 4E, bottom) or L655708 (Fig. 4F, bottom), consistent with previous observations of Duchen (1986) in mice, also studied in vitro.

In the awake rat (Fig. 4G), cutaneous nerve stimulation known to evoke PAD caused reflexes with repeated bursts of activity (Fig. 4H, top), again separated by about 7-10 ms (140 to 100 Hz), similar to that seen *in vitro* (Fig. E, left), suggesting that they may be due to the oscillating PAD-evoked spikes driving the oscillating reflexes. Indeed, hyperpolarizing the afferents with L655708 reduced these reflex oscillations (Fig. 4H, bottom) without changing the postsynaptic activity of the motoneurons assessed by the background EMG (Bkg, Fig. 4J), consistent with an action on the Ia afferents rather than on the motoneurons. L655708 reduced both the small monosynaptic reflex (# in Fig. 4H), as previously reported (Hari *et al*., 2021), and the repeated bursts of EMG at longer intervals. This is consistent with these long-latency reflexes being mediated by monosynaptic EPSPs triggered by PAD-evoked spikes, rather than polysynaptic reflexes from other sources, the latter which should be increased, rather than decreased, by reducing GABA action. Note that with L655708 the monosynaptic reflex (#) and some of the later putative PAD-evoked reflexes were facilitated with a prior brief cutaneous conditioning stimulation known to activate phasic PAD circuits and facilitate the monosynaptic EPSP by depolarizing the afferents (Fig. 4I), showing that the L655708 simply eliminated the EPSPs by hyperpolarizing the axons in Fig 4H, rather than indirectly acting to eliminate the EPSP by other pre- or postsynaptic pathways (e.g. via GABA_B_ receptors).

### Early soleus H-reflex inhibition following antagonist-evoked reflex associated with post activation depression

Similar to mice and rats, a conditioning electrical stimulation to the human flexor CPN supplying the TA muscle often produced an early excitatory reflex response in the extensor soleus muscle (early soleus reflex), which is not expected from our classical understanding of the reciprocal organization of reflexes, but is similar to what we observed with vibration, as detailed above. As described below, this early soleus reflex is consistent with a CPN-evoked PAD causing extensor afferent spikes and associated reflexes based on its latency and frequency of oscillation. Moreover, when we observed an early soleus reflex from antagonist CPN stimulation, subsequent soleus H-reflexes were suppressed (Figs. 5 & 6). In contrast, soleus H-reflexes were facilitated when we did not observe an early soleus reflex from the CPN stimulation (Fig. 7), indicating a non-spiking PAD that facilitated Ia conduction and motoneuron EPSPs, similar to that seen in rodents detailed above.

**Figure 5:**
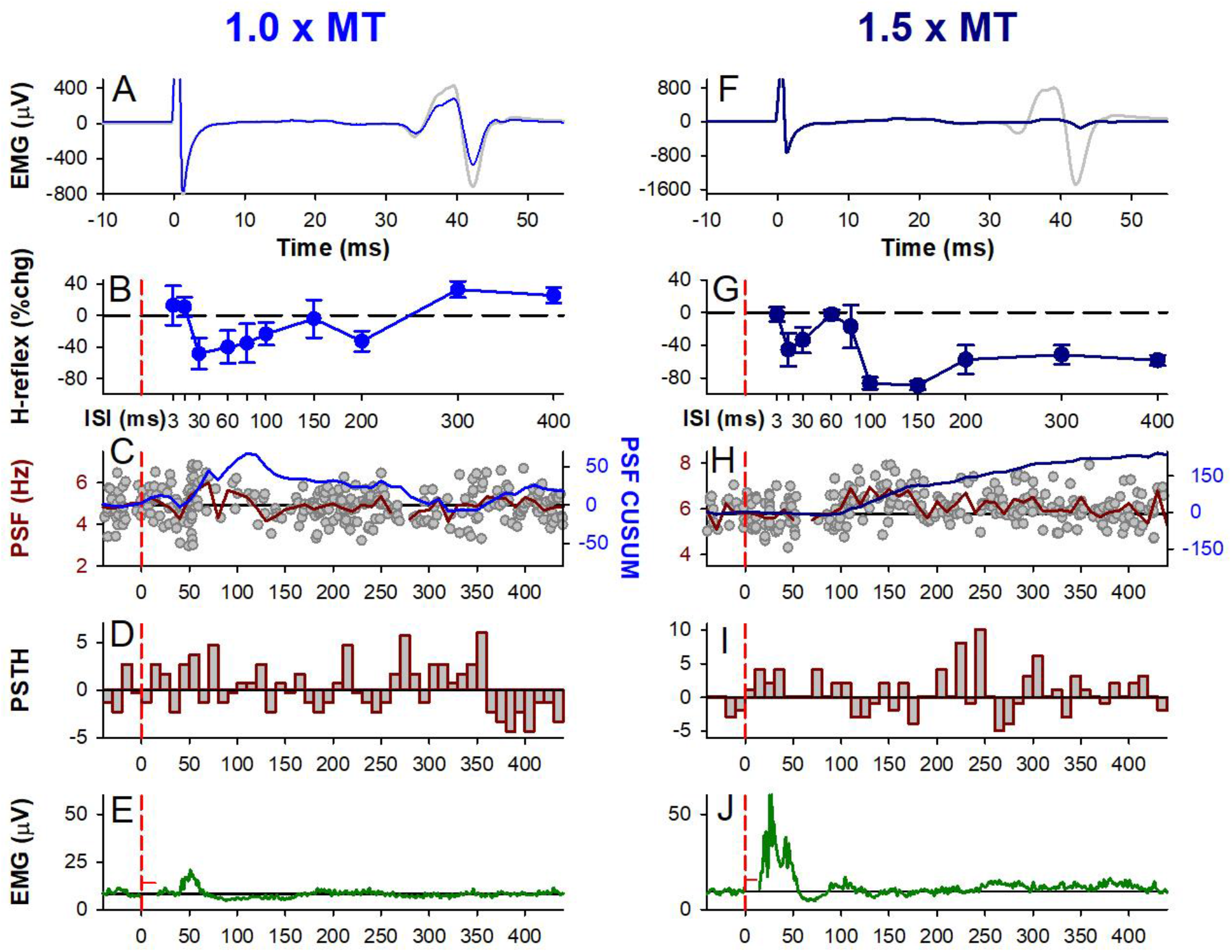
CPN stimulation: early H-reflex suppression. Same presentation as Figure 1 but for 1.0 x MT **(A-E)** and 1.5 x MT **(F-J)** electrical CPN conditioning stimulation, representative data from a single participant. **A,F)** Unrectified EMG of test (grey) and conditioned H-reflex (blue, 1.0 x MT at 30 ms ISI and dark blue, 1.5 x MT at 100 ms ISI). **B,G)** % change soleus H-reflex (Mean ± SD) plotted at each ISI tested. **C,H)** PSF (grey dots) with 108 sweeps in C and 98 sweeps in H, mean PSF (red line) and CUSUM of mean PSF (blue line). **D,I)** Number of motor unit counts per 10-ms time bin with average pre-stimulation count subtracted. **E,J)** Average rectified soleus EMG with 131 sweeps in E and 100 sweeps in J. Stimulation artifact has been removed (horizontal red line near 0 ms). **B-J)** Vertical dashed red lines indicate the onset of the CPN conditioning stimulation train (at time 0). * in J marks crosstalk from TA M-wave.

**Figure 6:**
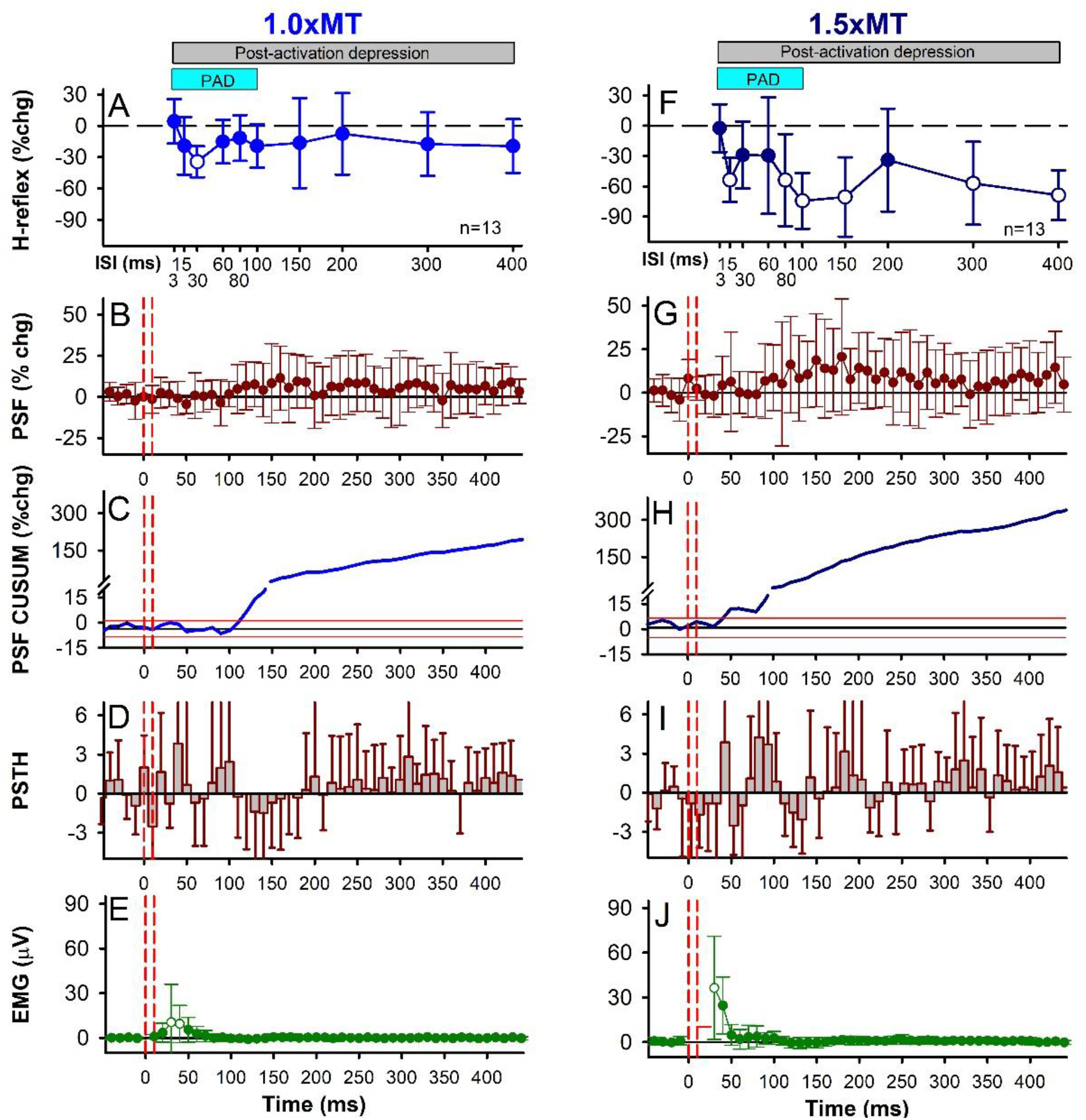
CPN stimulation: early H-reflex suppression, group data. Same presentation as Figure 2 but for the 1.0 x MT (A-E) and 1.5 x MT (F-J) CPN conditioning stimulation with data averaged across the 13 participants. The average size of the unconditioned H-reflex was 961.2 ± 454.1 μV (30.9±13.8% of Hmax; 16.6 ± 6.55% of Mmax). **A&F)** There was an effect of ISI on the % change of the H-reflex for both the 1.0 x MT (F[12,10] =2.392, *P* = 0.03) and 1.5 x MT (F[12,10] =7.801, *P* < 0.001, one-way RM ANOVAs) stimulus intensities, with the 30 ms ISI (*P* = 0.017, 1.0 x MT) and 15, 80, 100, 300 and 400 ms ISIs (P’s < 0.01, 1.5 x MT) significantly different from a 0% change (Tukey Test, white symbols). Blue bar indicates predicted duration of PAD and grey bar the predicted duration of post-activation depression. **B,G)** There was no effect of time on the % change PSF for the 1.0 x MT (F[12,61] = 1.094, *P* = 0.298) and 1.5 x MT (F[12,61] = 1.270, *P* = 0.087, CPN conditioning stimulation (one-way RM ANOVAs). **C,H)** % change CUSUM of the mean PSF averaged across participants. **D,I)** There was an effect of time on the count per bin (PSTH) for the 1.0 x MT (F[116,47] = 1.579, *P* = 0.010), but not the 1.5 x MT (F[116,47] = 1.352, *P* = 0.064), CPN conditioning stimulation (one-way RM ANOVAs). **E,J)** There was an effect of time on the EMG with the 1.0 x MT (F[12,46] = 3.361, *P* < 0.00) and 1.5 x MT (F[12,59] = 10.708, *P* < 0.001) CPN stimulation (one-way RM ANOVAs), with the 30 and 40 ms (P < 0.001, 1.0 x MT) and 30 ms (P < 0.001, 1.5 x MT) time bins significantly greater than 0 mV (Tukey Test, white symbols). Error bars ± SD.

**Figure 7:**
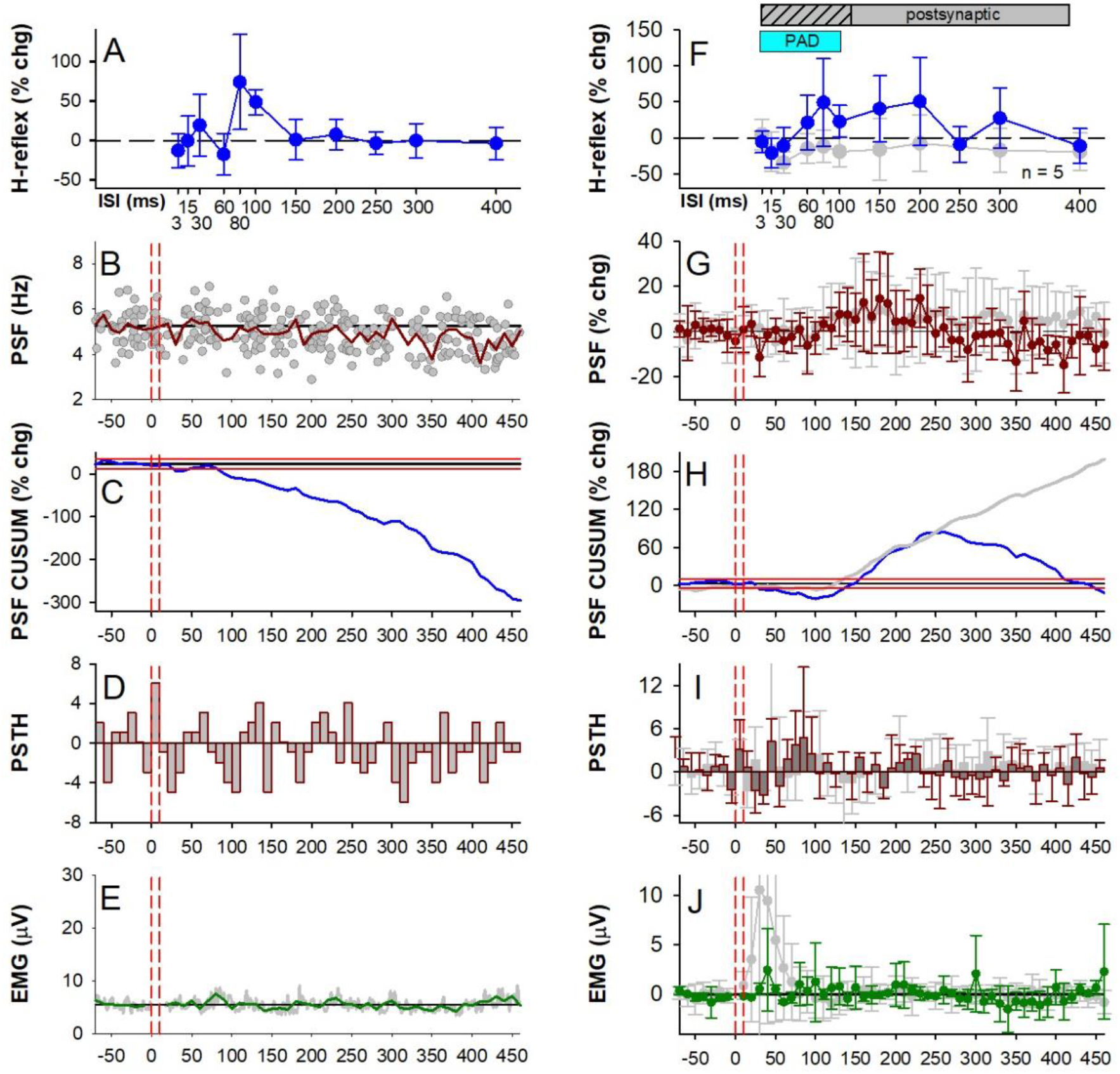
CPN stimulation: early H-reflex facilitation, individual and group data. Similar presentation as Figure 6 but with 1.0 x MT CPN conditioning stimulation that did not evoke an early reflex response in the SOL muscle. **A-E** data from an individual participant and **F-J** group data (n = 5 participants). Grey data points in F-J are replotted from Figs. 6A-E where 1.0 x MT CPN stimulation produced an early SOL reflex response. There was an effect of ISI on the % change H-reflex **(F)** (F[4,11] =2.113, *P* = 0.043) and time on the % change PSF **(G)** (F[3,61] = 1.673, *P* = 0.005) but no effect of time on the count per bin in the PSTH **(I)** (F[3,61] = 1.120, *P* = 0.281) or EMG **(J)** (F[3, 60] = 0.994, *P* = 0.498), all one-way RM ANOVAs. Error bars ± SD.

In participants where CPN conditioning produced an early soleus reflex that may be due to PAD evoked spikes (13/18 participants, Figs. 5 & 6), there was consistently an associated inhibition of the H-reflex. As shown for two of these participants, an early soleus reflex was readily apparent in the surface EMG near 40 ms at both CPN stimulation intensities (see arrows in Figs. 5E and J) that was more readily observed at 1.0 x MT since there was no crosstalk from the TA M-wave between 10 to 30 ms (* in Fig. 5J). Overall, the average amplitude of the early reflex response was larger for the 1.5 x MT stimulation (Fig. 6J) compared to the 1.0 x MT stimulation (Fig. 6E). This consistent reflex response produced marked synchronization in the firing probability of the soleus motor units, as shown in the PSTH (Figs. 6D,I; see also Fig. 5I) that was not readily apparent in the surface EMG. Interestingly, power spectrum analysis of the soleus EMG during the predicted early PAD window (i.e., 30-80 ms after the first CPN stimulation) revealed a frequency spike at 161.9 ± 27.4 Hz (not shown, 1.5 x MT), similar to the frequency of the PAD oscillations recorded in the VR in the rodent (140 Hz, Fig. 4 E).

The early soleus reflex was also reflected in the firing rate profiles of the soleus motor units in response to the CPN stimulation applied alone (PSF and PSF CUSUM), which reflect the profile of the underlying motoneuron membrane potential (Turker & Powers, 1999). The PSF increased early near 40 ms as shown for the 1.0 x MT stimulation in Figure 5C and in the group data where at the 1.5 x MT intensity, the PSF increased above 2 SD near 40 ms (Fig. 6H), which was around 7 ms later than the onset latency of the H-reflex (32.3 ± 2.0 ms) evoked from direct TN stimulation. Unlike the surface EMG, the PSF and PSF CUSUM displayed a longer-lasting increase following the CPN stimulation, lasting out to 400 ms as illustrated by the PSF remaining above the baseline rate (Fig. 6B,G) and by the sustained increase in the PSF CUSUM (Figs. 6C,H). This is consistent with multiple PAD-evoked Ia-EPSPs or a polysynaptic reflex response (see multiple Ia spikes in Figs. 3M and 4B-F). Similar effects are seen in cat motoneurons, where the flexor-evoked EPSPs appear in antagonist extensor motoneurons, sometimes alone (Frank, 1959) and sometimes riding on an IPSP (Stuart & Redman, 1992), and these too likely resulted from PAD-evoked spikes and caused post-activation depression of the Ia afferents (Hari et al. 2021).

Following the early soleus reflex, the inhibition of the H-reflex had two phases, an early phase starting at the 15-30 ms ISI (D1) and a later, more sustained phase of inhibition (D2) starting at the 100 ms ISI that was larger with the higher intensity 1.5 x MT stimulation (Fig. 5F,G) compared to the lower intensity 1.0 x MT stimulation (Fig. 5A,B), similar to previous studies (Mizuno *et al*., 1971; El-Tohamy & Sedgwick, 1983; Capaday *et al*., 1995). When averaged across the 13 participants, the soleus H-reflex was suppressed at the 30 ms ISI when conditioned with the 1.0 x MT CPN stimulation (Fig. 6A, white circle) and at several ISIs starting at 15 ms when conditioned with the 1.5 x MT CPN stimulation (Fig. 6F, see legend for statistics). In summary, the CPN conditioning stimulation produced an early and prolonged excitation in the soleus muscle indicative of PAD-evoked spikes in the soleus Ia afferent producing soleus monosynaptic EPSPs, as we have seen in rodents, and this EPSP activation of the soleus is associated with the long-lasting inhibition of the H-reflex, likely via post-activation depression of the H-reflex.

### Early soleus H-reflex facilitation in the absence of an antagonist-evoked reflex

Based on the findings in the mice, if a flexor conditioning stimulation does *not* activate a PAD-evoked spike in the extensor Ia afferents and thus, does *not* pre-activate the monosynaptic extensor EPSPs (as evident by a lack of early soleus reflex), then the H-reflex should not be inhibited by post activation depression, but if anything facilitated by PAD (Hari *et al*., 2021). Indeed, we found that when there was a lack of an early soleus reflex (and putative PAD evoked spikes) evoked by conditioning, the H-reflex was not inhibited, but instead facilitated by conditioning (seen in 5/18 participants in response to the 1.0 x MT CPN conditioning) [see also (Metz *et al*., 2021)]. This is shown for one participant in Figure 7 where the CPN stimulation does not increase the PSF or PSF CUSUM (Fig. 7B and C respectively) or the rectified surface EMG (Fig. 7E), whereas the H-reflex was facilitated at the 80 and 100 ms ISIs within the predicted PAD window. When averaged across these 5 participants, the H-reflex was facilitated between the 30 and 250 ms ISIs (blue trace, Fig. 7F) compared to the H-reflex suppression that occurred in the other group of 13 participants where an early reflex response was present (grey trace replotted from Fig. 6A). The amplitude of the H-reflex was significantly modulated across the different ISIs but post hoc analysis revealed that no H-reflex at a given ISI was different than a 0% change, likely due to the small number of participants [Fig. 7F, see legend for statistics]. However, this is consistent with our main conclusion that the lack of early reflex and associated PAD evoked spikes prevents an inhibition of the H-reflex, due to a lack of post activation depression. Further, when comparing the peak H-reflex facilitation that occurred at the 60 or 80 ms ISIs in each participant (during the predicted PAD window), the conditioned H-reflex was significantly greater than a 0% change (see also Fig. 3C in Metz *et al*., 2021). Note that on average, there was a brief period of motoneuron inhibition from 30 to 100 ms after the 1.0 x MT CPN stimulation, as reflected in the PSF (Fig. 7G), PSF CUSUM (Fig. 7H) and clustering of motor unit firing probability in the PSTH starting at 60 ms (Fig. 7I). This early motoneuron inhibition may have reduced the early facilitation of the conditioned H-reflex between the 15 to 60 ms ISIs within the PAD window. Following this early inhibition, there was a period of motoneuron facilitation as shown in the PSF (Fig. 7G, red) and PSF CUSUM (Fig. 7H, blue), but it was much briefer compared to when there was an early excitatory reflex response in the other group (the latter again plotted in grey from Fig. 6). Like in the PSF, the amplitude of the early excitatory reflex response in the soleus EMG was much smaller (or absent) in these 5 participants (Fig. 7J, compare green and grey traces).

As described above, our observed early excitatory reflex in the soleus EMG or PSF from the CPN conditioning stimulation may be indicative of PAD-evoked spiking in the soleus Ia afferents that produces long-lasting post-activation depression and subsequent H-reflex suppression. In contrast, when we observed no or very little early reflex activity is present, PAD evoked from the conditioning afferent input likely facilitates afferent conduction and thus increases subsequent H-reflexes. In further support of this, we observed that the amount of H-reflex inhibition at the 100 ms ISI was greater when the amplitude of the early excitatory reflex (measured between 30 - 50 ms) was also greater, as demonstrated by the negative correlation between these two measures (Fig. 8). Facilitation of the soleus H-reflex at the 100 ms ISI was typically produced only when there was no or very little early excitatory reflex activity in the soleus muscle

**Figure 8:**
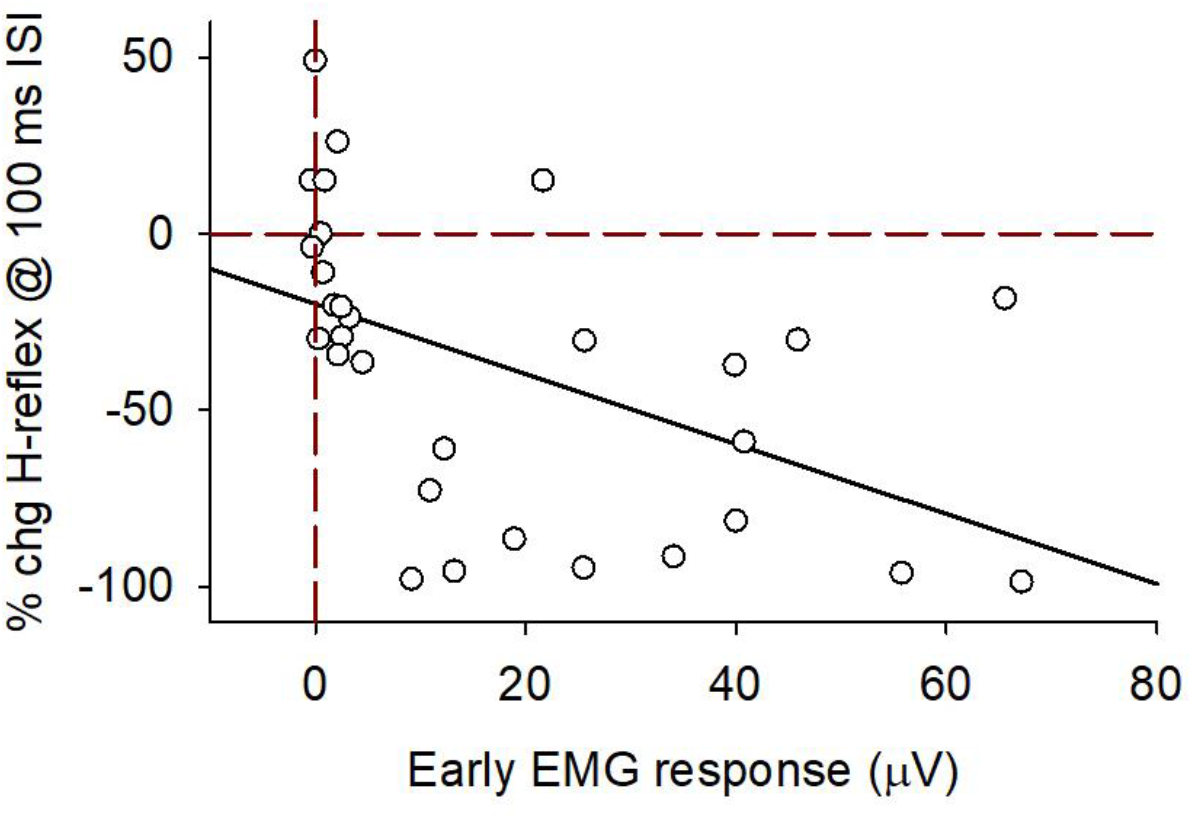
CPN stimulation: % change soleus H-reflex at the 100 ms ISI vs early soleus EMG response. % change (chg) H-reflex (100 ms ISI) plotted against the early EMG response (30-50 ms after CPN stimulation during slight contraction) for both the 1.0 and 1.5 x MT CPN stimulation for participants with and without an early soleus reflex (n = 18). Data fitted to a linear line (slope = 1.0 % chg /mV). There was a significant correlation between the % chg H-reflex and the early EMG response (r = −0.50, P = 0.005, n=30, Pearson Product-Moment Correlation). In some recordings (n = 6), the TA M-wave obscured the early soleus reflex and this data was not used. Red vertical dashed line marks no change in the soleus EMG from the CPN stimulation. Red horizontal dashed line marks 0 % change in the soleus H-reflex.

### Long lasting soleus H-reflex inhibition consistent with post activation depression

Given that conditioning of the flexor CPN may produce spikes in the extensor soleus Ia afferents and post activation depression of the H-reflex, we examined this further by exploring the *duration* of H-reflex inhibition by the CPN conditioning, which should be similar to the associated rate-dependent depression (RDD) of the H-reflex that is mediated by post activation depression, which can last seconds (Nielsen *et al*., 1993; Hultborn *et al*., 1996). Thus, we examined how long the H-reflex inhibition from the conditioning CPN stimulation lasted for in the same 13 participants that had the earlier suppression of the H-reflex presented above. As shown for two participants, the soleus H-reflex was maximally suppressed at the 500 ms ISI by a 1.5 x MT conditioning CPN stimulation and continued to be suppressed out to the 2500 ms ISI (Figs. 9Ai and Bi), which is similar in duration to the suppression of monosynaptic EPSPs seen in mice and rats when followed by a PAD-evoked spike and associated EPSP (Fig. 3E and Duchen, 1986). Across the group, the H-reflex was modulated at these very long ISIs, being significantly suppressed at all ISIs from 500 to 2500 ms (* Fig. 9Ci, see legend for statistics). By itself, the 1.5 x MT CPN stimulation increased the firing rate of the soleus motor units for at least 500 ms in some participants (PSF Fig. 9Bii) and for a briefer period in others (PSF Fig. 9Aii). Across the group, both the PSF (Fig. 9Cii) and the rectified surface EMG (Fig. 9Ciii) returned to pre-stimulus levels by 500 ms while the H-reflex continued to be suppressed (see statistics in legend). In contrast, the 1.0 x MT CPN stimulation intensity did not suppress the soleus H-reflex at these long ISIs as plotted for the 500 ms ISI (Fig. 9Di), consistent with it having a weaker or absent early soleus reflex that we take as evidence for PAD evoked spikes (Fig. 9Dii). Thus, the suppressed H-reflex at these long ISIs was not likely related to any direct post-synaptic inhibition of the soleus motoneuron at the time the H-reflex was evoked but rather, may have been produced by the earlier soleus reflex activation described above. In support of this, H-reflex suppression was significantly correlated to the early increase in PSF when data for the 1.0 and 1.5 x MT CPN stimulation intensities were plotted together (Fig. 9Diii) or when compared to the amplitude of the early EMG response to CPN stimulation (from 30 to 50 ms, r = −0.74, *P*<0.001, not shown).

**Figure 9:**
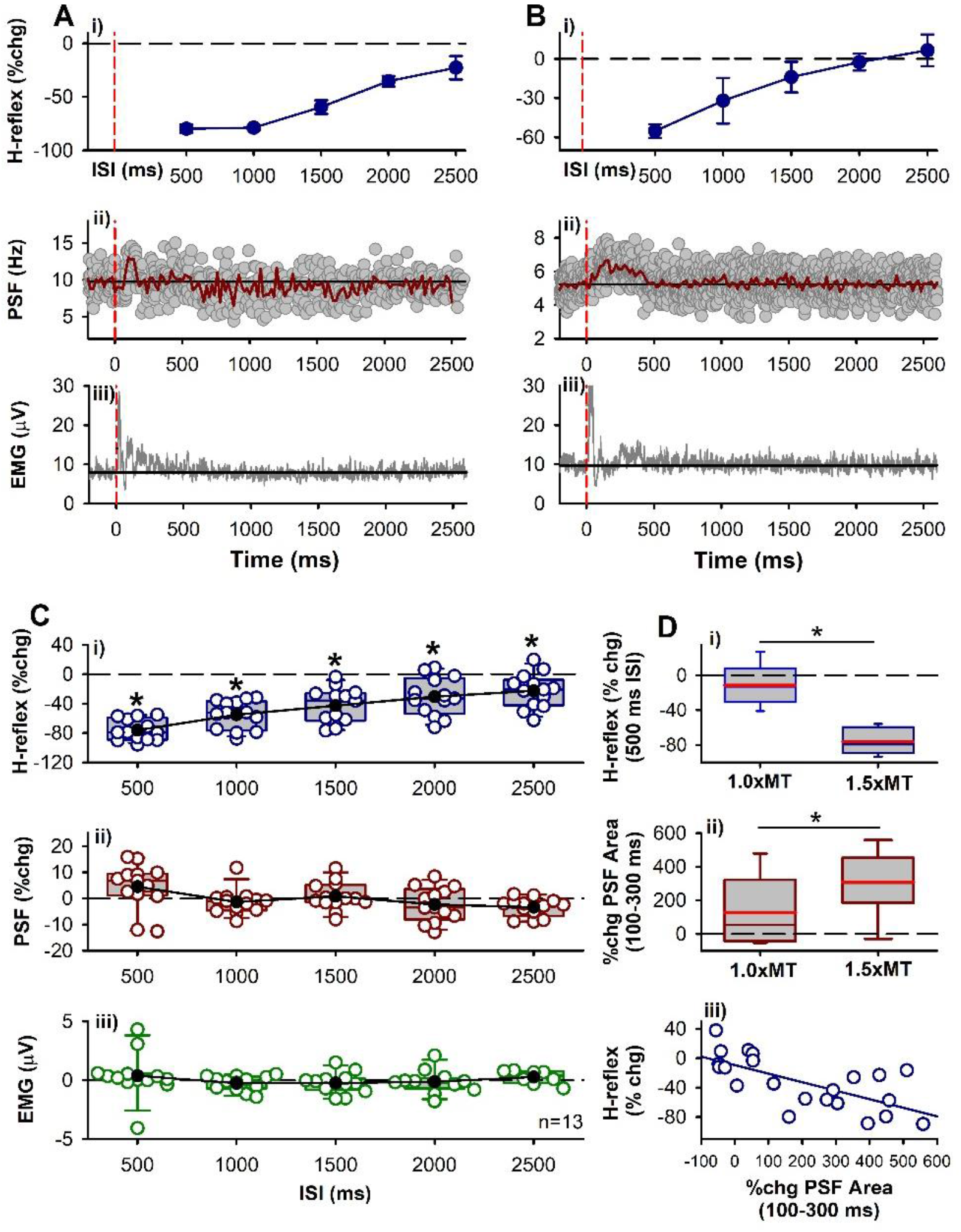
CPN stimulation: late H-reflex suppression. **A-B)** Representative data from two participants showing **i)** % change in H-reflex, **ii)** PSF (37 sweeps in A, 98 sweeps in B) and **iii)** rectified soleus EMG (40 sweeps in A, 30 sweeps in B). Vertical dashed red lines indicate time of 1.5 x MT CPN conditioning stimulation. **C)** Mean (black circles) and individual data points (open circles) from the 13 participants tested. There was an effect of ISI on the % change in the soleus H-reflex **(i)** (F[12,5] = 55.697, *P* < 0.001, one-way RM ANOVA) with all ISI’s significantly different from 0% change (*P* < 0.001, Tukey Test, marked by *). There was an effect of ISI on the % change PSF **(ii)** (F[12,5] = 4.393, *P* = 0.002) but not EMG (F[12,5] = 0.716, *P* = 0.614, one-way RM ANOVAs). The PSF and EMG data points occurred in the time bin that matched when the H-reflex was evoked at the various ISIs. **D)** Box plot (similar to Figs. 3C&G) of: **(i)** % change SOL H-reflex at the 500 ms ISI for the 1.0 and 1.5 x MT CPN conditioning stimulation, significantly different (P = 0.04, t-test); **(ii)**, % change in area of the PSF measured 100-300 ms after the 1.0 and 1.5 x MT CPN stimulation, significantly different (P < 0.001, t-test); **(iii)** % change H-reflex at 500 ms ISI plotted against % change PSF area (100 – 300 ms) with fitted linear line (blue, r = −0.7, P<0.001, Pearson Product-Moment Correlation). Error bars ± SD.

### Post-activation depression of the soleus H-reflex during RDD

We next investigated whether the early reflex activation in the soleus muscle by the antagonist CPN stimulation causes a similar long duration inhibition of the H-reflex to that expected from RDD, which has previously been demonstrated to be due to post activation depression, thus providing further evidence that PAD-evoked spikes in the soleus Ia afferents produces a long-lasting post-activation depression of the soleus H-reflex. That is, in 8 of the 13 participants from above, we examined if the profile of long-lasting suppression of the soleus H-reflex from antagonist CPN conditioning (CPN→TN) was similar to the profile of H-reflex suppression evoked by repetitive TN stimulation during RDD (TN→TN), the latter used to quantify the duration and profile of post-activation depression. Like with the CPN conditioning, repetitive stimulation of the TN inhibited the H-reflex at all intervals tested out to 2500 ms, with the greatest suppression at the 500 ms ISI (compare Figs. 10Ai & Aii). Across all participants, the average profile of H-reflex inhibition at the different ISIs was not different between antagonist CPN (dark blue) or homonymous TN (pink) conditioning stimulation trials (Fig. 10B, see legend for statistics), with both forms of inhibition lasting to 2500 ms. This suggested that the mechanism of H-reflex suppression was similar between the two stimulation protocols and likely due to post-activation depression.

**Figure 10:**
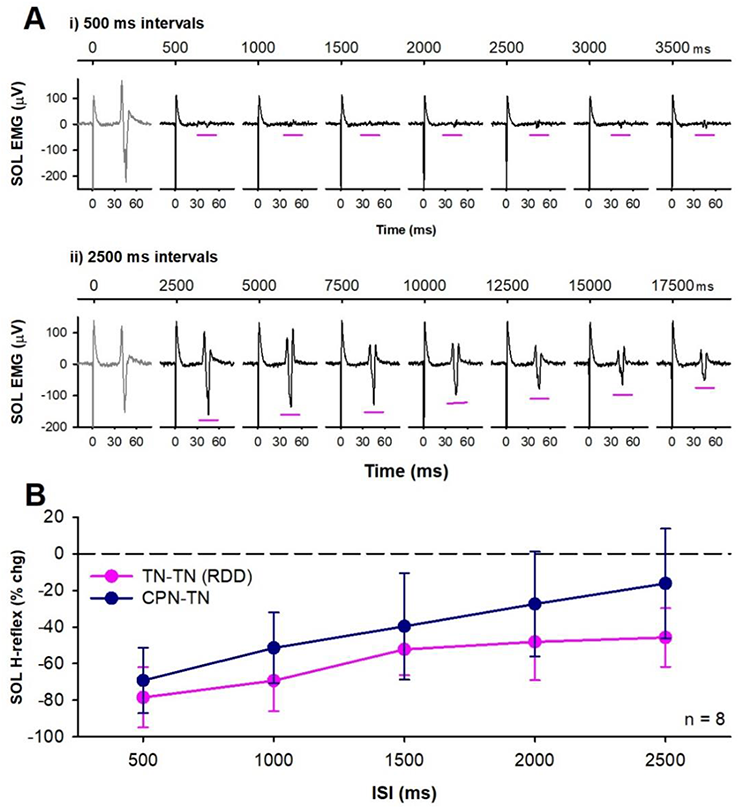
Post-activation depression of the soleus H-reflex. **A)** Example rate dependent depression (RDD) of unrectified soleus EMG evoked by repetitive TN stimulation every 500 ms (**i**) and 2500 ms (**ii**) in one participant. Continuous time displayed in top trace to show when TN stimulation was applied (marked by large stimulation artifact) and expanded time scale in bottom to display the H-reflex occurring in the time window marked by the pink horizontal lines (time reset to 0 ms when TN stimulation was applied). **B)** The % change in soleus H-reflex from repetitive TN stimulation (RDD, pink) averaged across the 8 participants at the different ISIs and % change H-reflex from CPN conditioning stimulation (dark blue), not significantly different (F[1,4] = 1.860, P = 0.152, two-way RM ANOVA). Error bars±SD.

## Discussion

Inhibitory control mechanisms within the spinal cord are needed to regulate the large influx of sensory information from the periphery that would otherwise exceed the computing capacity of the central nervous system (Rudomin & Schmidt, 1999). One way to reduce the flow of sensory information to spinal targets is through presynaptic inhibition of afferent terminals. To regulate the transmission of *proprioceptive* information, the depolarization of primary (Ia) afferents, or PAD, from the activation of GABA_A_ receptors was historically thought to inactivate sodium channels and/or shunt current at the afferent terminal and reduce neurotransmitter release to spinal motoneurons as one form of presynaptic inhibition (Willis, 2006). To demonstrate this, antagonist afferents were used to activate PAD in Ia afferents via GABAergic interneurons and shown to be accompanied by a parallel suppression of monosynaptic EPSPs in the motoneuron, though direct evidence for presynaptic inhibition via activation of terminal GABA_A_ receptors is lacking. For example, numerous studies have failed to find GABA_A_ receptors on Ia afferent terminals (Alvarez *et al*., 1996; Fink *et al*., 2014; Lucas-Osma *et al*., 2018; Hari *et al*., 2021). Instead these receptors are located more dorsally near the nodes of Ranvier near branch points where their activation facilitates afferent conduction and monosynaptic reflexes (Hari *et al*., 2021; Metz *et al*., 2021). We demonstrate here that the more likely mechanisms that produces the H-reflex suppression by antagonist afferents is both a direct long-lasting inhibition of the test motoneurons and an indirect post-activation depression of the Ia afferents mediating the H-reflex, two mechanisms that have been suggested previously but not tested explicitly (Eccles *et al*., 1961a; Hultborn *et al*., 1987a; Fink *et al*., 2014; Hari *et al*., 2021).

### Postsynaptic inhibition of motoneurons from antagonist afferents

We demonstrate here that the extensor H-reflex is inhibited for up to a half second following vibration of the flexor tendon, consistent with previous findings in cats (Curtis, 1998), which is too long to be explained by presynaptic inhibition from phasic PAD that only lasts for 100-200 ms, contrary to previous conclusions (Morin *et al*., 1984; Hultborn *et al*., 1987a; Hultborn *et al*., 1987b; Iles & Roberts, 1987; Roby-Brami & Bussel, 1990; Rossi *et al*., 1999). In humans, H-reflex suppression at ISIs of 40 to 500 ms was considered to be mediated primarily by PAD-induced presynaptic inhibition because it was thought that direct effects on the motoneuron from antagonist afferent conditioning did not last beyond the 40 ms ISI (Hultborn *et al*., 1987a; Hultborn *et al*., 1987b). However, as shown here and by others when using the PSF (Yavuz *et al*., 2018), inhibition of soleus motoneurons from a brief antagonist stimulation can last up to 300 ms. On average, the profile of PSF suppression from tendon vibration closely followed the early profile of H-reflex suppression following vibration, out to the 100 ms ISI (Fig. 2), and this motoneuron inhibition likely contributes to the suppression H-reflexes during this period. In cats and rodents, similar long-duration IPSPs and reduced neuronal time constants (τ, indicating postsynaptic shunting) have been induced in motoneurons from heteronymous afferent conditioning that also produced long-duration EPSP suppression (McCrea *et al*., 1990; Hari *et al*., 2021).

The prolonged inhibition of the soleus motoneurons may have resulted from a single long-duration IPSP (Pierce & Mendell, 1993; Hughes *et al*., 2005; Hari *et al*., 2021) or from multiple shorter-duration IPSPs triggered by multiple afferent inputs (Desmedt & Godaux, 1978). Regardless of the source, when using a weak conditioning stimulation to suppress H-reflexes like brief tendon vibration, sensitive methods are needed to measure if there are any direct actions on the motoneuron and for how long. The PSF provides a better representation of the IPSP evoked in the soleus motoneuron compared to the PSTH or surface EMG, the latter which better represent the occurrence of motoneuron discharge and are subject to synchronization artifact (Turker & Powers, 1999; Yavuz *et al*., 2018). In addition, we tested the excitability of the soleus motoneurons while they were tonically active during a weak voluntary contraction, so that any subtle inhibition from the tendon vibration or low intensity CPN stimulation could be revealed, in contrast to a resting motoneuron. A hyperpolarization of the distal dendrites by a low-intensity conditioning stimulation, which might not be detectible from intracellular somatic recordings in a resting motoneuron (McCrea et al., 1990), may produce a greater influence on the firing rate response of the cell (Hari et al, 2021). Thus, it is important to record single motor unit activity in a preactivated motoneuron to examine if the conditioning stimulation has any direct inhibitory effects on the motoneuron and the duration of its effect on the suppression of the H-reflex.

### Post-activation depression by antagonist afferents

Our finding that direct electrical stimulation to the flexor CPN, and sometimes even flexor tendon vibration, causes early extensor soleus reflexes led us to examine whether these unusual, excitatory reciprocal reflexes may contribute to subsequent post-activation depression of the soleus H-reflex. Consistent with previous findings, we found that H-reflexes are suppressed by the antagonist CPN conditioning (Mizuno *et al*., 1971; El-Tohamy & Sedgwick, 1983; Capaday *et al*., 1995). However, we demonstrate that this inhibition only occurred when short-latency reflexes to the soleus muscle are evoked by the conditioning electrical stimulation to the flexor CPN. The inhibition of the extensor H-reflex by electrical CPN stimulation occurred even though there was little evidence of postsynaptic inhibition of the soleus motoneurons compared to conditioning with tendon vibration. We propose here that the H-reflex suppression following this early reflex response is likely mediated by post activation depression of the soleus H-reflex induced by the antagonist CPN afferents activating PAD-evoked spikes in the soleus Ia afferents. In rodents we show that during a large and fast rising PAD, the membrane potential of the afferent can reach the sodium spiking threshold and produce action potentials that travel orthodromically down the Ia afferent to evoke monosynaptic EPSPs (early reflexes) in the motoneuron [see also (Duchen, 1986; Lucas-Osma *et al*., 2018)]. These monosynaptic EPSPs are likely produced by PAD-evoked spikes because: 1) the occurrence of the PAD-evoked spikes and the associated EPSPs are tightly linked in time with a monosynaptic latency [see also (Duchen, 1986)], 2) they repeat with a characteristic period of about the duration of the afferent AHP with a refractory period of 7 - 10 ms (140 – 100 Hz) in both rodent VR and human EMG recordings, and 3) both are reduced by specifically hyperpolarizing the afferent by blocking tonic PAD with L655708.

The PAD-evoked spike and the associated EPSP and excitatory soleus reflex likely cause post-activation depression that can have multiple underlying mechanisms, as detailed in the Introduction. These include the depletion of transmitter packaging and release from the Ia afferent, the afferent AHP and its refractory period (lasting up to 10 ms), and the indirect activation of GABA_B_ receptors on the afferent terminals triggered by the PAD-evoked spike to ultimately produce GABA_B_ mediated presynaptic inhibition (Fig. 11). Likely all these mechanisms are involved in the early inhibition of the extensor soleus H-reflex by flexor conditioning, but only the GABA_B_ action can account for the very long, up to 2 s, inhibition seen in our study and others (Eccles *et al*., 1961a; Curtis & Lacey, 1994, 1998) given that the recovery time constant from maximal transmitter depletion is around 300-400 ms (Neher & Sakaba, 2001). This raises the question of how are the GABA_B_ receptors on soleus afferent terminals activated by the flexor nerve stimulation? We do not favor the simple idea that flexor nerve stimulation directly activates the trisynaptic circuit that drives the GABAergic neurons that innervate the soleus afferent terminals and associated GABA_B_ receptors for two reasons. First, the inhibition of the soleus H-reflex or related mice EPSPs only occurs when it is linked to PAD-evoked spikes and associated PAD-evoked EPSPs or early soleus reflexes and thus, must be somehow indirectly linked to the GABA_A_ receptors that mediate PAD, as previously argued from the actions of GABA_A_ receptor blockers on this inhibition (Stuart & Redman, 1992). Second, the profile of inhibition from conditioning is similar in timing to RDD caused by repeated H-reflex activation, which has been shown to be mostly confined to inhibition mediated by the activation of the same afferents that are stimulated to evoke the H-reflex (thus also called homosynaptic depression) (Hultborn *et al*., 1996). This RDD is likely mediated by a restricted trisynaptic circuit, where the extensor afferent activates private GABAergic neurons that only activate extensor afferent terminals (Fig. 11). Thus, we favor the more complex scenario where flexor nerve stimulation evokes a widespread PAD and PAD-evoked spikes in extensor soleus afferents, which then activate a restricted trisynaptic circuit with GABAegic neurons that activate GABA_B_ receptors on the extensor afferent terminal (Fig. 11). This raises interesting questions about whether there are indeed subpopulations of different GABAergic neurons that innervate extensor vs flexor afferent terminals, which needs to be investigated in the future (Curtis & Eccles, 1960; Hultborn *et al*., 1996).

**Figure 11.**
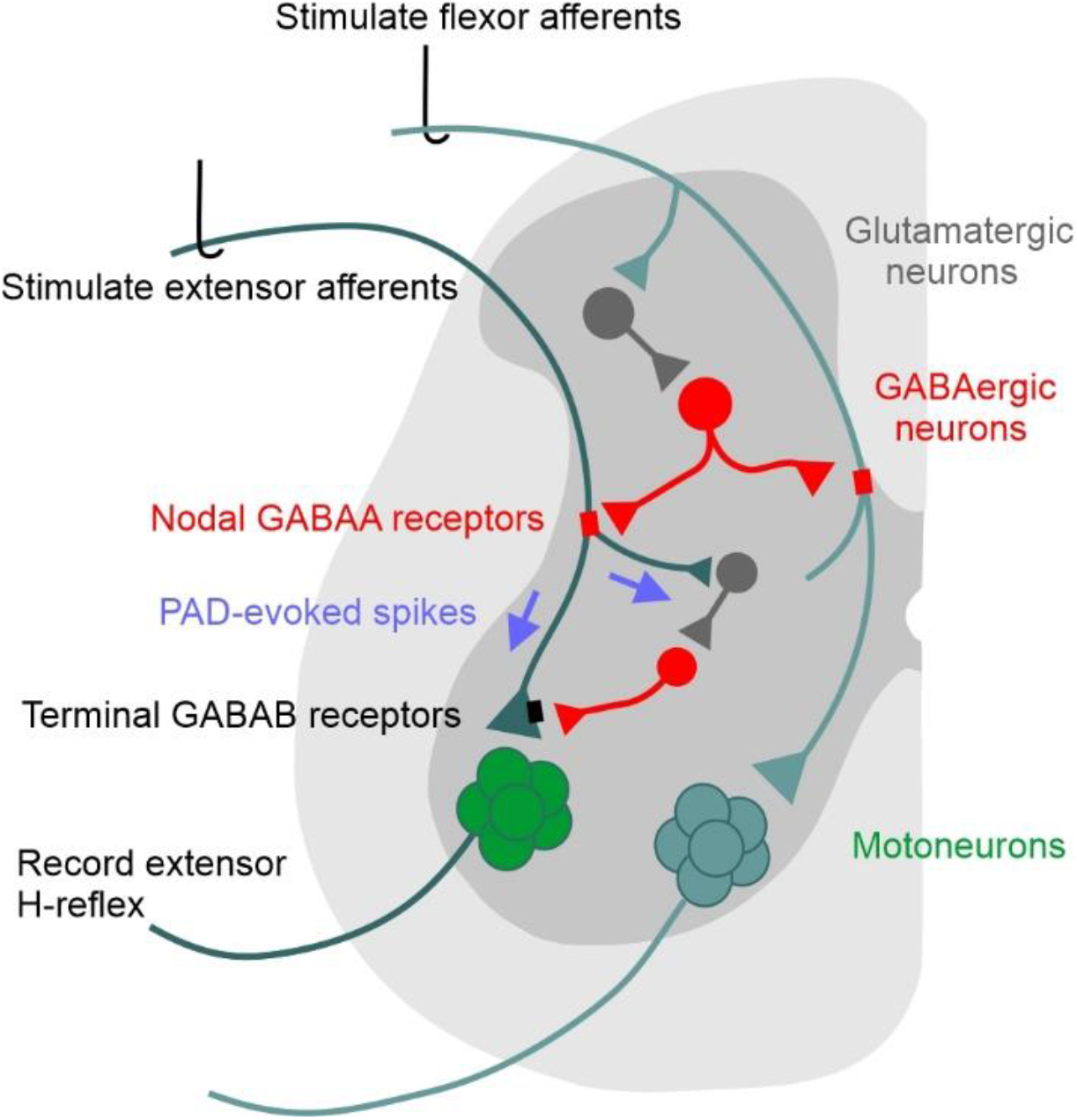
Schematic of PAD pathways mediating post-activation depression in the spinal cord. Activation of flexor afferents (cyan) activates a tri-synaptic dorsal PAD pathway (glutamatergic interneuron dark grey, GABA_axo_ neuron red) with axo-axonic projections to an extensor afferent (dark green), activating nodal GABA_A_ receptors (red) and PAD in the extensor afferent. If the resulting PAD in the extensor afferent reaches sodium spiking threshold, PAD-evoked spikes (purple arrows) travel orthodromically to: 1) depolarize extensor afferent terminals, evoking an EPSP in the extensor motoneurons and a subsequent MSR/H-reflex; 2) activate a trisynaptic circuit containing GABAergic neurons that activate GABA_B_ receptors (black) on the extensor afferent terminal. Thus, the PAD-evoked spike in the extensor Ia afferent can produce early and late post-activation depression of subsequently activated MSRs/H-reflexes by: 1) transmitter depletion following the PAD-evoked spike entering the extensor Ia afferent; 2) decreased afferent excitability from the post-spike refractory period that follows the PAD-evoked spike, and/or 3) inhibition of the afferent terminals by GABA_B_ receptors indirectly activated by the PAD-evoked spike (GABA_B_ mediated presynaptic inhibition).

For the flexor CPN to evoke PAD in extensor afferents, they must activate a circuit that that has GABAergic interneurons with axo-axonic projections (GABA_axo_ interneurons) onto extensor afferents, as just mentioned (Fig. 11). This arrangement is broadly consistent with our understanding of the widespread nature of PAD. That is, it has been known for many years in both animals (Eccles *et al*., 1961b; Willis, 1999) and humans (Shefner *et al*., 1992) that stimulation of afferents from one nerve can produce PAD and related PAD-evoked spikes (DRRs) in afferents from neighbouring nerves, since GABA_axo_ interneurons are innervated by first order neurons with projections across many segments within the spinal cord (Jankowska *et al*., 1981; Willis, 2006; Lucas-Osma *et al*., 2018). Furthermore, flexor nerve stimulation produces the strongest PAD in extensor afferents, compared to in flexor afferents (Eccles *et al*., 1961a; Rudomin & Schmidt, 1999). Here the new addition to this older concept is that PAD is likely mediated by GABA_A_ receptors at nodes, rather than ventral terminals (Hari *et al*., 2021), though this does not change the known distributions of PAD (Fig. 11). Thus, the amplitude or presence of the early reflex response in the soleus muscle evoked from the antagonist CPN stimulation can be used as a gross indicator of PAD-evoked spikes in the soleus Ia afferents. These Ia spikes likely produce post-activation depression in the soleus Ia afferent terminals to mediate the suppression of the conditioned H-reflexes, as we have discussed above. Indeed, the amplitude of the early soleus reflex activity is correlated with the amount of H-reflex suppression at the 100 ms ISI (Fig 8), an interval previously attributed to PAD-induced presynaptic inhibition of the Ia afferent terminal (Mizuno *et al*., 1971; Burke *et al*., 1992; Iles & Pisini, 1992; Capaday *et al*., 1995; Iles, 1996; Knikou & Mummidisetty, 2014; Howells *et al*., 2020). Moreover, when an early soleus reflex is not produced following low-intensity CPN stimulation, the conditioned H-reflexes are instead facilitated, likely due to a facilitation of spikes through the soleus Ia branch points by nodal PAD (Hari *et al*., 2021; Metz *et al*., 2021). Nodal depolarization and facilitation of Ia afferent conductance likely also occurs when H-reflexes were suppressed following the early soleus reflex and may account for the decrease in H-reflex suppression near the 30 to 60 ms ISIs during the rising phase of PAD that divide the D1 and D2 phases of inhibition (Mizuno *et al*., 1971; El-Tohamy & Sedgwick, 1983; Berardelli *et al*., 1987; Faist *et al*., 1996). However, this facilitation is probably masked by the stronger post activation depression of the Ia afferents from the PAD-evoked Ia spike(s).

Further evidence to support the conclusion that the profile of H-reflex inhibition from antagonist CPN conditioning is mediated by post-activation depression of the Ia afferent terminals is provided by its resemblance to the seconds long profile of rate dependent depression (RDD) from repeated TN stimulation that directly activates spikes in the soleus Ia afferents. Similar to RDD at long repetition intervals, H-reflexes are also suppressed out to 2.5 s after a strong conditioning CPN stimulation, well after any direct effect on the soleus motoneurons from the CPN stimulation had subsided. This is consistent with the long-duration suppression of the Ia-EPSP in mice and rats when a PAD-evoked Ia spike occurred 1500 ms earlier, even though the motoneuron had long returned to its resting membrane potential (Fig 3 and Duchen, 1986). Suppression of the H-reflex at these long intervals after conditioning occurrs only when there is an early reflex activation of the soleus motoneurons, as measured by both the PSF and rectified EMG. Alternatively, if this early-latency soleus reflex was solely mediated by a polysynaptic pathway from the CPN afferents that bypassed the soleus Ia afferents, the suppression of the H-reflex would likely not have lasted out to 2500 ms as we observe.

#### GABA_B_ receptor activation

Activation of GABA_B_ receptors on Ia afferent terminals by the flexor CPN stimulation may be involved in the suppression of the H-reflex and post activation depression as we discuss above, though we detail the mechanisms further here. Activation of GABA_B_ receptors produces presynaptic inhibition of Ia afferents and reflex transmission to motoneurons (Curtis & Lacey, 1994; Fink, 2013; Hari *et al*., 2021), likely by inhibiting voltage dependent calcium channels on the presynaptic boutons to reduce the calcium-dependent exocytosis of neurotransmitter following the arrival of an action potential at the afferent terminal (Curtis & Lacey, 1994; Curtis *et al*., 1997; Howell & Pugh, 2016). In contrast to GABA_A_ receptors, GABA_B_ receptors are densely located on Ia afferent terminals (Hari *et al*., 2021) and when activated by a brief train of conditioning afferent stimulation, they can suppress monosynaptic EPSPs for up to 800 ms (Curtis & Lacey, 1994, 1998) or more (Salio *et al*., 2017; Hari *et al*., 2021). This GABA_B_ mediated presynaptic inhibition is blocked by selective GABA_B_ receptor antagonists (Curtis & Lacey, 1994; Fink, 2013; Hari *et al*., 2021), with evidence for a spontaneous amount of steady presynaptic inhibition at rest (Hari *et al*., 2021), demonstrating how long it can last. Thus, GABA_B_ receptor activation on Ia afferent terminals could account for some of the long-lasting inhibition of the soleus H-reflex from the CPN conditioning stimulation or TA tendon vibration, requiring further study with specific antagonists to elucidate (Curtis *et al*., 1997). As mentioned above, we argue that these receptors are activated secondarily to PAD, with PAD-evoked spikes activating circuits that drive GABAergic innervation of extensor afferent terminals, as detailed in Fig. 11.

#### Passive PAD current to Ia terminals

Finally, although there are sparse GABA_A_ receptors at the afferent terminal, PAD from more proximal nodes may passively enter the afferent terminal to reduce the size of subsequent action potentials activated by the TN stimulation. However, because of the long distance of the last node from the Ia terminal (Hari *et al*., 2021) and the short space constant of the Ia axon, the amount of depolarization at the Ia afferent terminal is small (Lucas-Osma *et al*., 2018) and only reduces the size of the action potential by 1% even when nearby. This reduction is not likely to have a noticeable effect on neurotransmitter release and suppression of H-reflexes (Hari et al., 2021).

### Conclusion and clinical implications

We have provided several lines of evidence that H-reflex suppression from antagonist afferents is produced, in part, by direct effects on the motoneuron and from post activation depression of Ia afferent transmission, the latter mediated by PAD-evoked Ia spikes triggered from the conditioning afferent input. These findings bring into question the interpretation of many human experiments claiming that H-reflex suppression by antagonist/heteronymous afferents are produced by PAD-mediated presynaptic inhibition of the Ia afferent terminal [reviewed in (Stein, 1995; Misiaszek, 2003; Hultborn, 2006; Willis, 2006)]. By extension, the idea that presynaptic inhibition of Ia afferents from PAD is reduced following nervous system injury or disease also needs to be reexamined, with examples including spinal cord injury (Ashby & Verrier, 1975; Mailis & Ashby, 1990; Roby-Brami & Bussel, 1990; Azouvi *et al*., 1993; Calancie *et al*., 1993; Faist *et al*., 1994; Aymard *et al*., 2000; Knikou & Mummidisetty, 2014; Caron *et al*., 2020), cerebral palsy (Mizuno *et al*., 1971; Achache *et al*., 2010), brain injury (Koelman *et al*., 1993; Faist *et al*., 1994), stroke (Milanov, 1992; Koelman *et al*., 1993) and multiple sclerosis (Azouvi *et al*., 1993; Koelman *et al*., 1993; Nielsen *et al*., 1995). Because the amount of H-reflex suppression has been incorrectly equated to the amplitude of PAD, changes in the activation of GABA_A_-receptor mediated PAD and its role in the regulation of afferent conduction in these various disorders needs to be reexamined. This is important because drug therapies that enhance GABA_A_ receptor activation, such as the benzodiazepine diazepam (Valium), have been used to treat spasticity in some of these conditions (Simon & Yelnik, 2010; Chang *et al*., 2013). The greater use of GABA_B_ antagonists, such as baclofen, to treat spasticity aligns better with our new understanding of the role of GABA_B_ in presynaptic inhibition and post activation depression. Interestingly, diazepam increases PAD measured in the dorsal part of the afferent and also reduces the monosynaptic reflex (Stratten & Barnes, 1971). Thus, drug therapies that enhance the activation of GABA_A_ receptors could actually increase afferent transmission while paradoxically increasing post-activation depression through the triggering of PAD-evoked Ia spikes or by directly facilitating inhibitory GABA_A_ receptors on the motoneuron (Cartlidge *et* al., 1974; Simon & Yelnik, 2010; Chang *et al*., 2013). In light of our new findings, we need to investigate how PAD evoked in dorsal parts of the Ia afferent is involved in the abnormal control of afferent conduction and transmission following injury or disease of the central nervous system in order to develop better methods to improve spared sensorimotor function and treat spasticity.

## Notes

### Competing Interest Statement

The authors have declared no competing interest.

